# Local retinoic acid directs emergence of the extraocular muscle functional unit

**DOI:** 10.1101/2020.01.07.897694

**Authors:** Glenda Comai, Marketa Tesarova, Valerie Dupé, Muriel Rhinn, Pedro Vallecillo Garcia, Fabio da Silva, Betty Feret, Katherine Exelby, Pascal Dollé, Leif Carlsson, Brian Pryce, Francois Spitz, Sigmar Stricker, Tomas Zikmund, Jozef Kaiser, James Briscoe, Andreas Schedl, Norbert B. Ghyselinck, Ronen Schweitzer, Shahragim Tajbakhsh

## Abstract

Coordinated development of muscles, tendons, and their attachment sites ensures emergence of functional musculoskeletal units that are adapted to diverse anatomical demands among different species. How these different tissues are patterned and functionally assembled during embryogenesis is poorly understood. Here, we investigated the morphogenesis of extraocular muscles (EOMs), an evolutionary conserved cranial muscle group that is crucial for the coordinated movement of the eyeballs and for visual acuity. By means of lineage analysis, we redefined the cellular origins of periocular connective tissues interacting with the EOMs, which do not arise exclusively from neural crest mesenchyme as previously thought. Using 3D imaging approaches, we established an integrative blueprint for the EOM functional unit. By doing so, we identified a developmental time window where individual EOMs emerge from a unique muscle anlage and establish insertions in the sclera, which sets these muscles apart from classical muscle-to-bone type of insertions. Further, we demonstrate that the eyeballs are a source of diffusible retinoic acid that allow their targeting by the EOMs in a temporal and dose dependent manner. Using genetically modified mice and inhibitor treatments, we find that endogenous local variations in the concentration of retinoids contribute to the establishment of tendon condensations and attachment sites that precede the initiation of muscle patterning. Collectively, our results highlight how global and site-specific programs are deployed for the assembly of muscle functional units with precise definition of muscle shapes and topographical wiring of their tendon attachments.

## INTRODUCTION

Acquisition of shape and pattern during development depends on the orchestrated crosstalk between a variety of tissues and cell types. While significant knowledge on the mechanisms of differentiation and patterning within individual tissues has been attained, much less is known on how patterning of different adjacent tissues is integrated. The vertebrate musculoskeletal system serves as an ideal model to study these processes as different tissues including muscle, tendon and their attachments need to be articulated in three-dimensions (3D) for proper function (Hasson, 2011; Huang, 2017).

Among the craniofacial muscles, the morphological configuration of the extraocular muscles (EOMs) has been a longstanding challenge in comparative anatomy and evolutionary biology. Besides specialized adaptations, the basic EOM pattern is shared among all vertebrate classes (Noden and Francis-West, 2006; Suzuki et al., 2016; Young, 2008) and includes 4 recti muscles (the superior rectus, the medial rectus, the inferior rectus, and the lateral rectus) and two oblique muscles (superior oblique and inferior oblique) for movement of the eyeball. Most vertebrates have also accessory ocular muscles involved in protecting the cornea (Noden and Francis-West, 2006; Spencer and Porter, 2006): the retractor bulbi, which serves to retract the eye in several groups of vertebrates and the levator palpabrae superioris, which is not directly associated with the eyeball but controls eyelid elevation in mammalian species. As such, the EOMs constitute an archetypal and autonomous functional unit for the study of how muscles, tendons and tendon attachments are integrated with the development of the eyeball, their target organ.

Craniofacial muscles are derived from cranial paraxial and prechordal head mesoderm (Noden and Francis-West, 2006; Ziermann et al., 2018) and the corresponding connective tissues, i.e. tendons, bones and cartilages and muscle connective tissue surrounding myofibres, were reported to be derived from cranial neural crest cells (NCC) (Nassari et al., 2017; Ziermann et al., 2018). Although early myogenesis is NCC-independent, NCC later regulate the differentiation and segregation of muscle precursors, dictate the pattern of muscle fiber alignment, and that of associated skeletal and tendon structures (Bohnsack et al., 2011; Ericsson et al., 2004; Noden, 1983b; Rinon et al., 2007; Tokita and Schneider, 2009; von Scheven et al., 2006). Moreover, studies of mouse mutants in which several genes were deleted in NCC cells have explicitly demonstrated their non-cell autonomous role in muscle morphogenesis at the level of the jaw (McGurk et al., 2017; Rinon et al., 2007; Shimizu et al., 2018), extraocular (Evans and Gage, 2005; Heude et al., 2015) and somitic-derived tongue muscles (Iwata et al., 2013). However, the full series of events driving morphogenesis of craniofacial musculoskeletal functional units is to date unexplored, probably due to the anatomical complexity of their configuration in the head. Understanding the developmental mechanisms that allow musculoskeletal connectivity is essential to understand their anatomical diversification during the evolution of the vertebrate head. Yet, proximate factors that allow the coordinated emergence of the individual muscle masses with that of their tendons and attachment sites are poorly defined.

To-date, much of the work on the tissue interdependency during musculosketal development has been done in the developing limb. Lateral plate mesodem derived connective tissue cells and tendon primordia establish a pre-pattern and restrict the locations where limb muscle progenitors proliferate and differentiate (Kardon, 1998; Kardon et al., 2003) prior to segregation (splitting) into individual muscle masses by several extrinsically controlled mechanisms (Hasson, 2011; Sefton and Kardon, 2019). Limb tendons are initially established independently of muscle, but their later development is modular, and depend on muscle or cartilage according to their location in the limb (Huang, 2017). Finally, generation of bone superstructures enable anchoring of muscles via tendons to the skeleton. Bone superstructures are initiated independently of muscle, by a unique set of progenitors that co-express SOX9 and Scleraxis (Scx) (Blitz et al., 2013; Blitz et al., 2009; Sugimoto et al., 2013), but remarkably, muscle and superstructure patterning seem to be regulated in a coordinated manner (Colasanto et al., 2016; Swinehart et al., 2013). However, given the striking differences in gene regulatory networks governing cranial muscle development (Diogo et al., 2015; Sambasivan et al., 2011), as well as the diverse embryological origins of connective tissue cells in the head (Nassari et al., 2017; Ziermann et al., 2018), it is unclear if this established logic of musculoskeletal integration is conserved in the head and for muscles that do not operate with muscle-to-bone attachments such as the EOMs.

It has been proposed that the developing central nervous system, sensory organs, pharyngeal endoderm and surface ectoderm provide spatial patterning cues to cranial connective tissue precursors (Noden and Francis-West, 2006; Ziermann et al., 2018). In this context, all-trans retinoic acid (ATRA), the biologically active metabolite of retinol (vitamin A), is a critical morphogen with widespread roles in craniofacial development (Gitton et al., 2010; Rhinn and Dolle, 2012). ATRA acts as a ligand for nuclear retinoic acid receptors (RARs), which are ligand-dependent transcriptional regulators that work as heterodimers with retinoid X receptors (RXRs) (Cunningham and Duester, 2015; Rhinn and Dolle, 2012). ATRA is synthetized from retinol through two oxidation steps by specific retinol/alcohol dehydrogenases (RDH/ADH) and retinaldehyde dehydrogenases (ALDH1A1, ALDH1A2 and ALDH1A3) and catabolized into hydroxylated derivatives by enzymes of the cytochrome P450 subfamily (CYP26s) (Cunningham and Duester, 2015; Rhinn and Dolle, 2012). ATRA was shown to be critical for early eye development in several species (Bohnsack et al., 2012; Harding and Moosajee, 2019; See et al., 2008), where the enzymes responsible of ATRA metabolism are expressed in the early retina with tight spatiotemporal patterns (Cvekl and Wang, 2009). Several studies have demostrated that through production of RA, the developing eye acts as a signaling center nucleating anterior segment morphogenesis, with the transcription factor *Pitx2* as the potential major downstream effector in periocular NCC (Bohnsack et al., 2012; Matt et al., 2005; Matt et al., 2008; Molotkov et al., 2006). However, whether ATRA is required for morphogenesis of the adjacent EOMs and associated connective tissues remains unexplored.

Here, by a combination of 3D-mesoscopic imaging approaches, we present the first integrative blueprint for morphogenesis of the EOM functional unit. We provide genetic evidence for the existence of a retinoic acid signaling module that coordinates the emergence of individual EOMs, their tendons and insertion sites. We show that the action of retinoic acid signaling in muscle patterning is mainly non-cell autonomous, through its action on the neural crest-derived periocular mesenchyme. Furthermore, we characterise the dynamics of EOM-tendon insertion sites, which are impaired upon ATRA-deficiency. Overall, our results show that in spite of the distinct embryological origins of these tissues in the head, interactions between muscles, tendons and their attachments are similar to those observed in the limb, yet they exhibit specifc hallmarks that are characteristic of this anatomical location. This includes regionalized apoptotic foci that presage tendon attachment sites in a context that is distinct from the canonical tendon to bone configuration.

## Results

### Genetic fate mapping of mouse periocular tissues

To investigate the morphogenesis of the EOMs in the mouse, we focused first on adjacent tissues whose development should be coordinated with that of the EOMs. The periocular mesenchyme (POM) is a heterogeneous cell population surrounding the optic cup that gives rise to specialized structures of the anterior segment of the eye and connective tissues associated with the EOMs (Williams and Bohnsack, 2015). With exception of the EOMs and endothelial lining of ocular blood vessels (choroid), all connective tissues of the POM (cartilage, muscle connective tissue, tendons), were reported to be derived from NCC in zebrafish, chicken and mouse embryos (Couly et al., 1992; Creuzet et al., 2005; Gage et al., 2005; Grenier et al., 2009; Johnston et al., 1979; Noden, 1983a). However, the extent of the POM in relation to its different cell populations, has not been fully assessed. Thus, we investigated the expression pattern of PITX2, a well established marker of both the POM and EOM progenitors (Gage et al., 2005), *Scx*, a tendon progenitor marker (Schweitzer et al., 2001) and SOX9, which is expressed in NCC and required for determination of the chondrogenic lineage (Mori-Akiyama et al., 2003). We simultaneously traced the contribution of different embryonic populations using *Mesp1^Cre^;R26^tdTomato^* and *Wnt1^Cre^;R26^tdTomato^* mice that label the mesodermal and NCC derivatives (Danielian et al., 1998; Saga et al., 1999) in combination with the *Tg:Scx-GFP* transgenic line (Pryce et al., 2007).

Coronal sections at the level of the eye at E13.5 showed that *Scx-GFP*, SOX9 and PITX2 expression delimited a triangular area of cells medial to the eyeball encompasing the EOMs (dotted lines, Fig 1A-D), however, with variable intensities along the medio-lateral axis. SOX9 and PITX2 were more strongly expressed laterally between the retina and ectoderm, where the EOM tendons insert. The *Scx-GFP* reporter labelled the future EOM tendons and also muscle connective tissue as described for other muscles (Deries et al., 2010). As expected, EOM tendons and muscle connective tissue at the insertion level were of NCC origin (*Wnt1^Cre^;R26^tdTomato^*, Fig 1A-D). Surprisingly, we found that lineage contributions differed in dorsal sections. The future Annulus of Zinn, the tendinous ring from which the recti muscles originate at the base of the skull (Spencer and Porter, 2006), as well as the muscle connective tissue at this level, were of mesodermal origin (*Mesp1^Cre^;R26^tdTomato^*, Fig 1E-H), although they also co-expressed *Scx-GFP*, SOX9 and PITX2 at this stage. The dual mesoderm/NCC contribution to POM connective tissues was observed from E11.5 (not shown) to E17.5 (Suppl Fig 1A-F). These findings indicate that the EOMs develop in close association with connective tissues of two distinct embryonic origins: neural crest laterally (at the EOM insertion level) and mesoderm medially (at the EOM origin level).

**Fig1.**
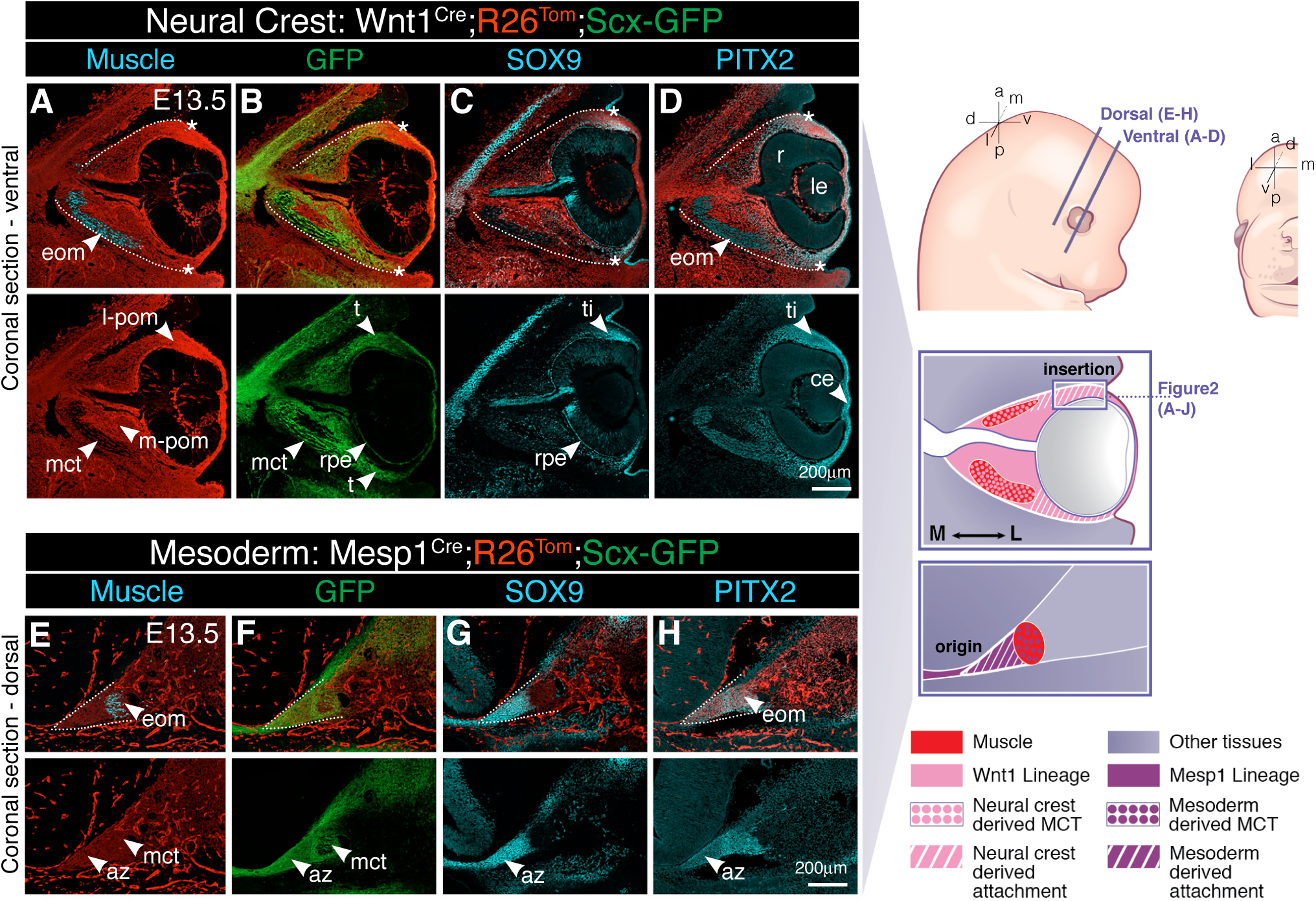
Lineage contributions to the EOM functional unit. **(A-H)** Neural Crest cell (NCC, *Wnt1^Cre^;R26^Tom^*) and mesoderm (*Mesp1^Cre^;R26^Tom^*) lineage contributions to the periocular region on coronal cryosections of E13.5 embryos, combined with immunostaining for GFP (*Tg:Scx-GFP* tendon reporter), muscle (PAX7/MYOD/MYOG, myogenic markers), SOX9 (chondrogenic/scleral marker) and PITX2 (EOM muscle progenitor and periocular mesenchyme marker). Sections at ventral (A-D) and dorsal (E-H) levels. Asterisks in 1A-D denote tendon insertion sites. Dashed regions in A-H delimitate the periocular mesenchyme. Panels A-B and E-F, an anti-DsRed antibody was used to enhance the R26^Tom^ signal. Panels C-D and G-H, the endogenous reporter signal was used. C,D and G,H are consecutive sections to the ones shown in panels A,B and E,F respectively. (n=3 per condition). az, annulus of zinn; ce, corneal ectoderm; eom, extraocular muscles; le, lens; l-pom, lateral periocular mesenchyme, at the level of the tendon insertions; mct, muscle connective tissue; m-pom, medial periocular mesenchyme; r, retina; rpe, retinal pigmented epithelium; s, sclera; t, tendon (at the level of the insertion); ti, tendon insertion site.

### Development of the EOMs and their insertions overlap spatio-temporally

Patterning of the EOMs and their insertions has been largely disregarded due to their complex anatomy, which is difficult to interpret from tissue sections alone. Therefore, we established an imaging pipeline that includes whole mount immunofluorescence (WMIF) of the periocular region, tissue clearing, confocal microscopy and reconstructions of the obtained images into 3D objects. To visualize the developing EOM we used MYOD/MYOG/Desmin as myogenic commitment and differentiation markers and MF20 to label myofibers (Fig 2A-C’, Video 1). At E11.75, the EOMs were present as a single anlage (Fig 2A-A’) medial to the eyeball. By E12.5, the EOM anlage split towards the POM-ring into the submasses corresponding to the future four recti, two oblique muscles and the accesory retractor bulbi muscle (Fig 2B-B’). Fully separated muscle subsets were evident by E13.5 (Fig 2C-C’). Thus, we conclude that EOM patterning in the mouse occurs by splitting from a single anlage with major morphogenetic changes taking place between E11.75 and E12.5.

**Fig2.**
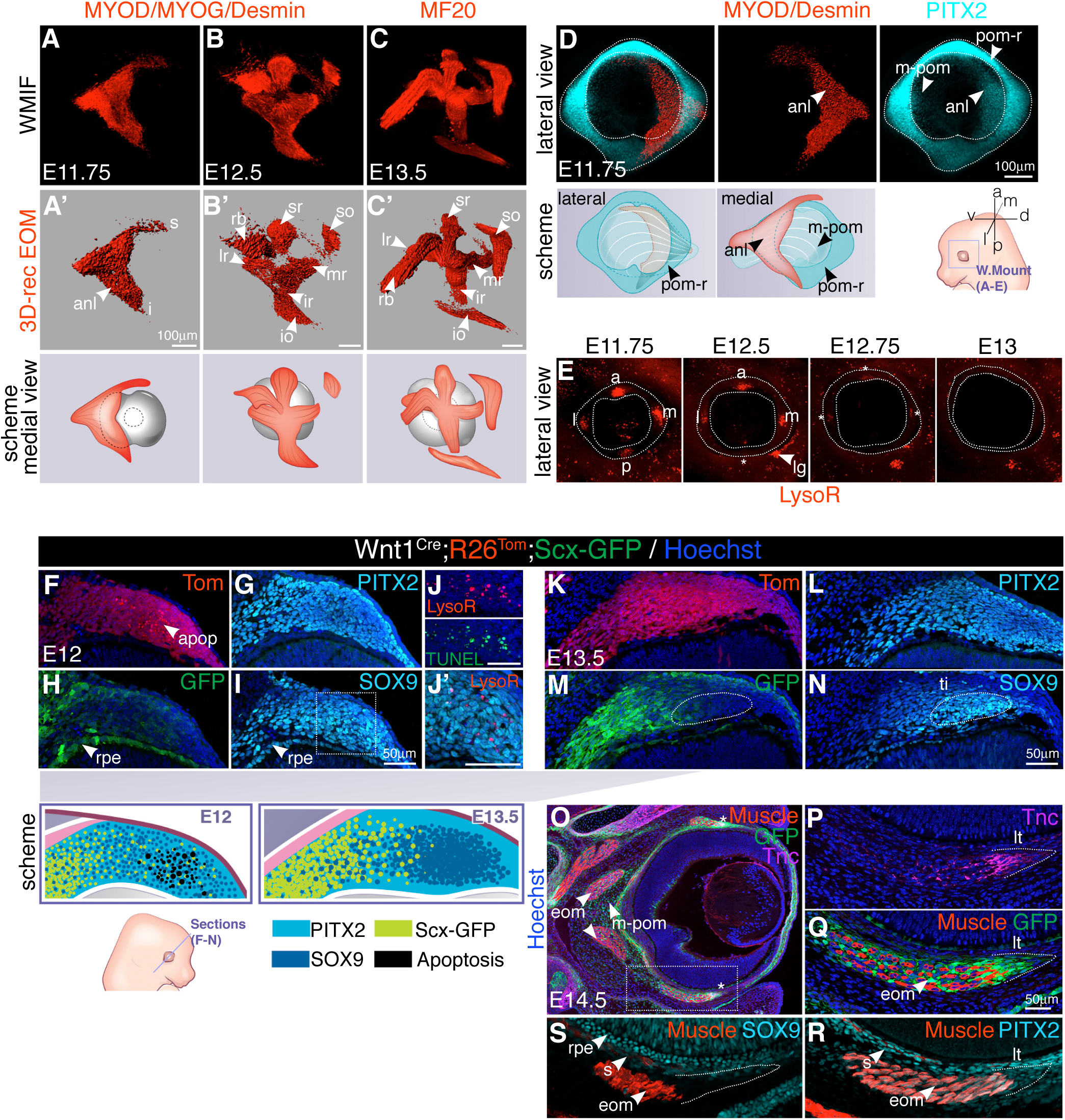
Developmental timing of EOM functional unit components. **(A-C)** WMIF for the MYOD/MYOG/Desmin (myogenic differentiation markers) (A, B) and MF20 (myofibers) (C) at the indicated embryonic stages. Medial views (left eye). **(D)** WMIF for MYOD/Desmin (labeling the EOM anlage, shown as isosurface for clarity) and PITX2 (labelling the POM and EOM anlage) on E11.75 embryos (left eye). Lateral and medial views as schemes. **(E)** Whole Mount LysotrackerRed staining of the periocular region at the indicated stages. The POM is delimited with a dotted line. Anterior (a), posterior (p), medial (m) and lateral (l) apototic foci are indicated. Asterisks indicate apoptotic foci with reduced intensity from E12.5 onwards. **(F-I)** Immunostaining of lateral POM in coronal sections of E12 *Wnt1^Cre^;R26^Tom^;Scx-GFP* embryos. Images correspond to insertion site of superior rectus muscle in POM. G, immunostaining of section adjacent to the one shown in (F,H,I; channels split for clarity). **(J,J’)** Tunel, LysotrackerRed and SOX9 staining of area depicted in I. Scale bars 50 microns. **(K,N)** Immunostaining of lateral POM in coronal sections of E13.5 *Wnt1^Cre^;R26^Tom^;Scx-GFP* embryos. Images correspond to insertion site of superior rectus muscle in POM. L, immunostaining of section adjacent to the one shown in (K,M,N; channels split for clarity). **(O-S)** Immunostaining on coronal sections of E14.5 *Scx-GFP* embryos. Tnc (Tenascin), GFP and PITX2 co-localize in tendons at level of insertion. MF20 (O-Q) and Smooth muscle actin (SMA) (R,S) were used to label EOM muscle. Dashes in R,S were drawn according to GFP labelling on adjacent sections. Asterisks in O denote tendon insertion sites. Low magnification views of panels P-S are shown in Suppl Fig 2. (n=3 per condition). anl, anlage; apop, apoptosis spots; eom, extraocular muscles; ir, inferior rectus; le, lens; lg, lacrimal gland sulcus; t, tendon (at the level of the insertion); lr, lateral rectus; i, inferior EOM anlage projection; io, inferior oblique; mr, medial rectus; pom-r, periocular mesenchyme ring; m-pom, medial periocular mesenchyme; rpe, retinal pigmented epithelium; s, sclera; s, superior EOM anlage projection; sr, superior rectus; so, superior oblique; rb, retractor bulbi.

The EOMs insert into the sclera, a dense fibrous layer derived from the POM. Foci of cell death in the POM were suggested to label the tendon attachment positions of the 4 recti muscles (Sulik et al., 2001). Thus, to better understand the establishment of the EOM insertions in the POM within the time window of EOM patterning, we performed whole mount immunostaining for the POM marker PITX2 and LysoTracker Red staining to detect programmed cell death on live embryos (Fogel et al., 2012) (Fig 2D,E). WMIF for the PITX2 at E11.75 revealed a ring of PITX2^high^ cells immediately adjacent to the retinal pigmented

epithelium (RPE) (POM-ring) that formed a continuum with PITX2^low^ cells extending towards the base of the EOM anlage (medial-POM) (Fig 2D). Lysotracker Red staining revealed a highly dynamic pattern in the POM-ring, where four foci were present at E11.75 at the horizons of the eye, which regressed from E12.5 onwards (Fig 2E). To confirm that the apoptotic foci define tendon attachment positions in the POM, we performed whole mount imunostaining for EOM progenitors and GFP (*Tg:SCX-GFP*) on Lysotracker stained embryos. Surprisingly, we observed that even before muscle splitting initiated, *Scx-GFP+* condensations bridged the edges of the EOM anlage and the four Lysotracker+ foci in the POM (E11.75, Video 2), presaging the locales of the future individual muscles (E12.5, Video 3).

To examine the EOM insertions with higher resolution, we immunostained *Wnt1^Cre^;R26^tdTomato^*;*Scx-GFP* coronal sections with POM markers (Fig 2F-N). At E12, PITX2 and SOX9 showed high expression in the lateral POM at the insertion site, while *Scx-GFP* was mostly expressed in a salt and pepper pattern at this level (Fig 2G-I). Where cells appeared more compact, as assessed by SOX9 and PITX2 staining, the Tomato staining appeared punctate (Fig 2F), reminiscent of apoptotic cells. Cell death was confirmed by LysoTracker Red and TUNEL staining on adjacent sections (Fig 2J, J’). By E13.5, PITX2 continued to mark a large periocular region, but SOX9 and *Scx-GFP* expression patterns were segregated and apoptotic foci undetectable (Fig 2K-N). Sox9 was expressed more laterally in a condensed group of cells probably representing the lateral tendon attachment points (Fig 2M), preceded medially by strong *Scx-GFP* staining at the tendon tips (Fig 2N). By E14.5, lateral EOM tendons expressed *Scx-GFP*, Tenascin and also PITX2 (Fig 2P-R, Suppl Fig 2A-H). At this stage, expression of SOX9 and PITX2 were donwregulated in the POM and reduced to the thin scleral layer (Fig 2R, S, Suppl Fig 2E-L). Altogether, these results show that patterning of the EOMs, their tendons and insertion sites overlap spatio-temporally and thus might be regulated in a coordinated manner. Moreover, similarly to other locations in the body, *Scx* and SOX9 show complementary patterns at the EOM insertion site, but additional specifc hallmarks, notably the presence of apoptotic foci and PITX2 expression, seem to be characteristic of this site.

### Abnormal EOM morphogenesis in mutants with ocular malformations

To obtain further insights into the development of the EOM functional unit, we next focused on the role of the target organ, the eyeball, on EOM patterning. To this end, we performed micro-computed tomography (micro-CT) scans in mouse mutants with a spectrum of ocular perturbations. First, we analyzed *Pax6^Sey/Sey^* embryos, which are anophthalmic where eye development is arrested at the optic vesicle stage (Grindley et al., 1995; Kaufman et al., 1995; Roberts, 1967). In heterozygote embryos, EOM patterning proceeded as in wildtype controls, where 4 recti and 2 oblique muscles were identifiable by E13.5 (Fig 3A, B). In contrast, *Pax6^Sey/Sey^* mutant EOMs appeared as a unique sheet of tissue on top of a rudimentary optic vesicle (Fig 3C). Given that Pax6 is expressed in the optic vesicle, giving rise to the retina and RPE, as well as in the overlying ectoderm that forms the lens and cornea (Grindley et al., 1995), but not in the EOMs, these observations suggest a non-cell autonomous role in EOM patterning. Similarly, in *Lhx2^Cre^;Lhx2^fl/fl^* embryos, where inactivation of the *Lhx2* gene in eye committed progenitor cells leds to a degeneration of the optic vesicle at E11.5 (Hagglund et al., 2011), EOM patterning was severely affected and few EOM submasses could be observed (Suppl Fig 3A-B’). Finally, in cyclopic embryos resulting from inversion of Shh regulatory regions, EOMs underwent splitting and projected towards the centrally located ectopic eye, although with an abnormal 3D arrangement as reported for human cyclopia conditions (Suppl Fig 3C-C’’) (Ziermann et al., 2018). Together with previous studies where surgical removal of the developing eye results in smaller EOMs with limited spatial organization (Bohnsack et al., 2011; Twitty, 1932; von Scheven et al., 2006), our observations reinforce the notion that the eye is not only a target organ for muscle insertion, but it plays a critical role as an organizer of EOM patterning. However, as severe eye defects are associated with abnormal migration of NCC to the periocular region (Langenberg et al., 2008; Matsuo et al., 1993), it is difficult to pinpoint the underlying cause of the observed EOM phenotypes. Thus, we decided to assess directly whether the developing eye could act as a source of molecular signals to induce the geometrically complex EOM pattern and that of their associated connective tissues.

**Fig3.**
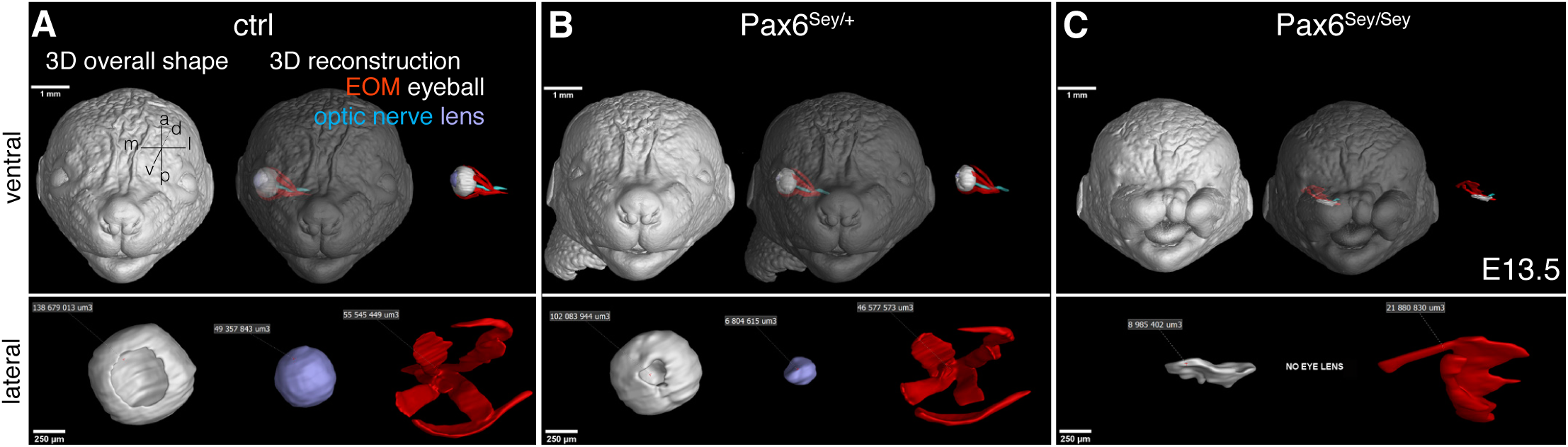
EOM morphogenesis in mutants with eye defects. **(A-C)** µCT-based 3D reconstruction of EOM, eyeball, optic nerve and lens in E13.5 control (A), *Pax6^Sey/+^* (B) and *Pax6^Sey/Sey^* embryos (C). Note that in heterozygote embryos EOM patterning proceeds normally despite having a smaller retina and lens (n=3).

### Muscle patterning depends on retinoic acid signaling of neural origin

To study the role of target organ derived cues in EOM patterning, we investigated the role of retinoic acid signaling, which plays multiple paracrine roles during embryonic eye development (Cvekl and Wang, 2009). As ALDH1A1-3 are the rate-limiting enzymes in the production of ATRA (Duester, 2009), we characterised their expression in the periocular region at the time of EOM patterning. Between E10.5 and E12.5, *Aldh1a1* was expressed strongly in the dorsal retina, lens and corneal ectoderm (Suppl Fig 4A), *Aldh1a2* was expressed in the temporal mesenchyme (Suppl Fig 4A-A”), and *Aldh1a3* was mostly restricted to the ventral retina and corneal ectoderm (Fig 4A, Suppl Fig 4A) (Matt et al., 2005; Molotkov et al., 2006). Interestingly, ALDH1A3 was the only retinaldehyde dehydrogenase protein detected in the RPE and optic stalk/optic nerve between E10.5-E12.5 and thus centrally positioned with respect to EOM development (Fig 4A). To target the retinoic signaling pathway (Fig 4B), we used *Aldh1a3^-/-^* (Dupe et al., 2003) and *Rdh10^-/-^* (Rhinn et al., 2011) mutants. We also administered the BMS493 inhibitor (pan-RAR inverse agonist) to pregnant females every 10-12h between E10.5-E11.75, i.e. preceeding the initiation of muscle splitting (Fig 2A-C) and once NCC migration to the POM has finalized (Osumi-Yamashita et al., 1994; Serbedzija et al., 1992). Micro-CT analysis showed that *Aldh1a3^-/-^* and BMS493-treated embryos displayed eye ventralization and a shortened optic nerve when compared to controls (Suppl Fig 4B), but they retained the overall organization of the nasal capsule and orbit (Suppl Fig 4C). Ventral and lateral views of the 3D-reconstructed EOMs (Fig 4C-E), showed that *Aldh1a3^-/-^* and BMS493-treated embryos lacked the standard 3D arrangement of 4 recti and 2 oblique muscles observed at E13.5 in control embryos. Nevertheless, in all cases EOMs originated medially from the hypochiasmatic cartilages of the presphenoid bone, indicating that the overall orientation of the EOMs was preserved (Fig 4D’’’). Thus, given that the EOMs are more affected at their insertion than their origin level upon ATRA deficiency, this finding suggests that EOM patterning is in part modular.

**Fig4.**
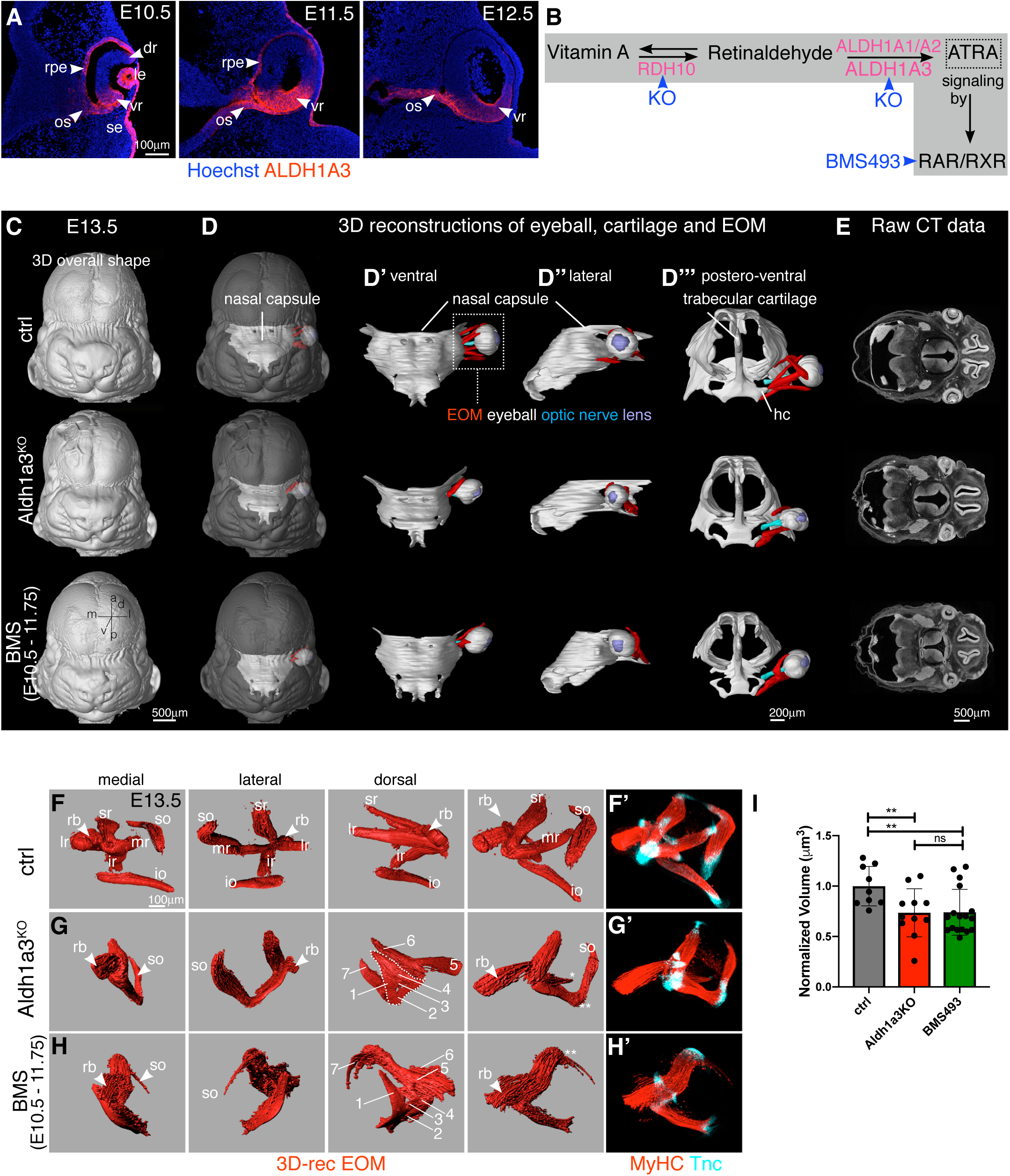
Extraocular muscle morphogenesis is dependent on ATRA. **(A)** Immunostaining for ALDH1A3 on coronal sections of E10.5, E11.5 and E12.5 control embryos (n=2). **(B)** Scheme of retinoic acid signaling pathway with key enzymes for oxidation of retinol and retinaldehyde (pink) and mutants/inhibitors used in this study (blue). **(C-D’’’)** µCT-based 3D-reconstruction of chondrogenic mesenchymal condensations of nasal capsule, trabecular cartilage, EOM, eyeball, optic nerve and lens in E13.5 control, *Aldh1a3^KO^* and BMS treated embryos. EOM visualization in context of whole head (D), nasal capsule (D’-D’’), trabecular capsule (D’’’). **(E)** Raw µCT data of panels in C-D (n=2) **(F-H)** EOM 3D-reconstructions of whole mount MyHC immunostaining of E13.5 control (F), *Aldh1a3^KO^* (G) and BMS treated embryos (H) (n>9). (1-7) denote non-segregated muscle masses with differential fiber orientation (see also Suppl Fig 5E). Raw immunostaining data for MyHC (muscle) and Tenascin (Tnc, tendon) are shown in panels F’-H’. **(I)** Relative EOM volume (compared to control) of E13.5 3D-reconstructed whole mount immunostained EOM of E13.5 control, *Aldh1a3^KO^* and BMS treated embryos. Each dot represents an individual embryo (n>9). dr, dorsal retina; hc, hypochiasmatic cartilage; io, inferior oblique; ir, inferior rectus; le, lens; lr, lateral rectus; mr, medial rectus; os, optic stalk; se, surface ectoderm; so, superior oblique; sr, superior rectus; rb, retractor bulbi; rpe, retinal pigmented epithelium; vr, ventral retina

To analyze EOM and tendon patterning with higher resolution, we performed whole mount immunostainings for differentiated myofibres (MyHC) and tendon (Tenascin) (Fig 4F-H’). On medial and lateral views of 3D-reconstructed EOMs, only the retractor bulbi and superior rectus could be clearly identified among the non-segregated muscle fibres in *Aldh1a3^-/-^* embryos (Fig 4G, G’). As expected from global invalidation of retinoic acid signaling, EOM perturbation was more severe in BMS493-treated embryos (Fig 4H, H’). The SO was absent, or continuous with the anterior part of the anlage, and the medial portion of the retractor bulbi was thicker and less clearly isolated from the rest of the anlage. In both conditions, Tenascin and *Scx-GFP+* cells were present at the tips of the individual, though mispatterned, muscles at this stage (Fig 4G’,H’, Video 4). Analysis of *Aldh1a3^-/-^* and BMS493-treated embryos revealed on average a 26% reduction in the EOM volume compared to controls (Fig 4I). However, MyHC+ myofibres were present in *Aldh1a3^-/-^* and BMS493-treated embryos (Fig 4G,H), suggesting that EOM differentiation was not overtly affected in these conditions. Instead, these observations suggest that EOM fiber alignment and segregation is dependent on retinoic acid signaling.

As dose and temporal control are critical in the context of retinoic acid signaling (Chawla et al., 2016; Gitton et al., 2010; See et al., 2008), we performed other BMS493 injection regimes between E10.5 and E12.5 (Table 1). EOM patterning phenotypes categorised as strong or severe at E13.5 were only obtained when a E10.75 time point of injection (8PM of day E10.5) was included in the regime (experiment type I-III, Table 1, Suppl Fig 4D). Surprisingly, even a single injection at this time point resulted in strong phenotypes (experiment type IV, Table 1, Suppl Fig 4D), while exclusion of this timepoint (experiment type VI and V, Table 1, Suppl Fig 4D) resulted only in mild phenotypes at best. Therefore, we identified an early and restricted temporal window where ATRA activity, prior to any sign of muscle splitting, impacts correct muscle patterning 36-48 hours later. Given the critical action of RDH10 in generating retinaldehyde, the intermediate metabolite in the biosynthesis of ATRA, we examined EOM development in *Rdh10* mutant embryos. *Rdh10* is normally expressed in the optic vesicle and RPE (Sandell et al., 2007), and in *Rdh10^-/-^* embryos, eyes develop intracranially, close to the diencephalon with a very short ventral retina (Rhinn et al., 2011). WMIF analysis of *Rdh10* mutant EOMs revealed that the anlage was specified, but lacked any sign of segmentation, in agreement with an upstream role in ATRA synthesis (Suppl Fig 4E,F). Taken together, our data show that retinoic acid signaling is essential for the correct myofiber alignment and segregation of EOM masses in a dose and time dependent manner.

**Table 1.**
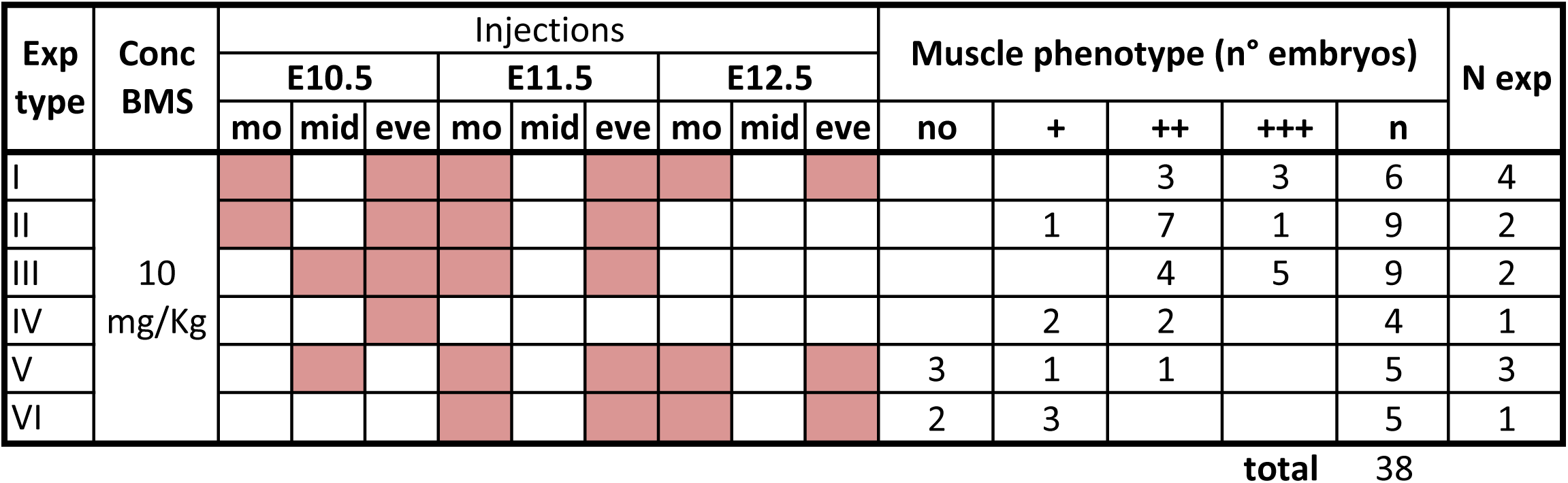
BMS493 injection regimes. The same BMS493 concentration (Conc BMS) was used across injection regimes, from I to VI. Pink cases mark injection time points in the morning (mo), midday (mid) or evening (eve) between E10.5 and E12.5. The number of embryos with different muscle phenotype severity at E13.5 are shown: (no), no altered phenotype; mild (+), muscle mispatterning but overall organization retained; strong (++), strong mispatterning but with a minumum of 2 muscles split; severe (+++), no or almost absent splitting. n is the total number of embryos analyzed for a certain treatment type (n=38 embryos analyzed in total); N exp, number of times experiment was repeated.

### Muscle patterning is dependent on ATRA-responsive cells in the periocular mesenchyme

To track cells that are responsive to ATRA and responsible of EOM patterning we used a novel retinoic acid transgenic reporter line (Schedl et al., unpublished) that comprises three *RARE* elements from the *Rarb* gene fused to the *Hsp68* promoter driving expression of a tamoxifen-inducible *Cre-ERT2* recombinase. By crossing this line with the *R26^mTmG^* reporter mice (Muzumdar et al., 2007) (Fig 5A, *Rare^CreERT2^;R26^mTmG^*), we permanently labelled ATRA-responsive cells and their descendants with membrane GFP expression. We found a greater amount of responsive cells in the periocular region when Tamoxifen was administered between E9.75-E10.5 (Suppl Fig 5A-C). To assess if myogenic cells and the adjacent POM respond to retinoic acid signaling, we scored the co-expression of GFP (reporter) and myogenic markers (PAX7, MYOD, MYOG) in bulk cell preparations from the periocular region. Notably, the great majority of reporter positive-responsive cells were not myogenic and thus were part of periocular conective tissues (88% if induction at E9.75; 98% if induction at E10.5) (Fig 5B,C). Upon induction of the reporter at E9.75, and analysis on tissue sections and by WMIF at E11.75 and E12.5, we observed that the mesenchyme around the optic nerve in close proximity to the developing EOMs was positive for the reporter (Fig 5D-E’,H). Interestingly, the overal number of POM GFP+ cells increased from E11.75 to E12, which is the temporal window corresponding to EOM splitting (Fig 5D-E’). The SOX9+ POM-ring that includes the EOM attachments also had abundant numbers of GFP+ cells (Fig 5F-G’,H’). Finally, inactivation of downstream retinoic acid signaling with BMS493, prior and subsequent to the induction of *Rare^CreERT2^;R26^mTmG^* mice with Tamoxifen (Suppl Fig 5D), greatly reduced the responsiveness of the reporter in the POM compared to non-treated controls and led to muscle mispatterning (Suppl Fig 5E-H, Videos 5 and 6). Together, these results are in agreement with previous studies (Matt et al., 2005; Matt et al., 2008; Molotkov et al., 2006) showing that despite a sophisticated pattern of ATRA metabolism in the developing retina, retinoic acid signaling exerts its action mostly non-cell autonomously, in the adjacent POM, and within a temporal window that is crucial for EOM patterning.

**Fig5.**
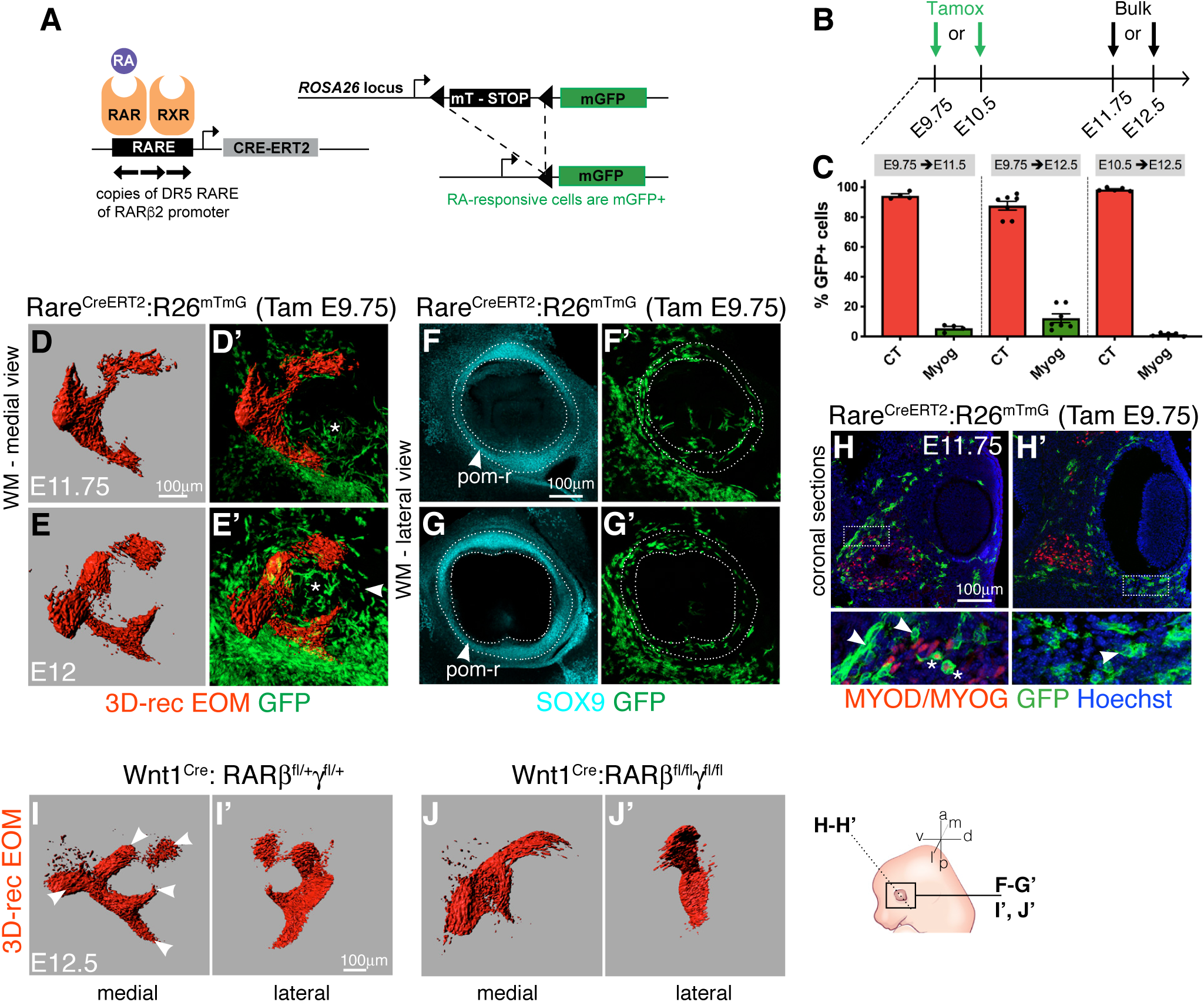
Periocular connective tissues are responsive to retinoic acid signaling. **(A)** Scheme of mouse alleles used. *Tg:RARE^CreERT2^* line is composed of three RARE elements from the *Rarb* gene fused to hsp68 promoter driving expression of a tamoxifen-inducible *Cre^ERT2^* recombinase. Crossing this line with the *R26^mTmG^* reporter line, allows permanent labelling of the ATRA-responsive cells and tracing of their descendents. **(B)** Strategy used to determine cell-types reponsive to retinoic acid signaling in *Tg:RARE^CreERT2^::R26^mTmG^* embryos. Tamoxifen was injected to pregnant females at E9.75 or E10.5 and cells of the periocular region isolated at E11.75 or E12.5. **(C)** Percentage of recombined (GFP+) cells in connective tissue (CT) and myogenic (Myog) populations for different treatments. Each dot represents an individual embryo (n>4 embryos/condition). **(D-G’)** Whole mount immunostaining for SMA (smooth muscle actin, differentiated muscle), GFP (ATRA-responsive cells) and SOX9 (periocular mesenchyme) at indicated embryonic stages. (D-E’) 3D-reconstructed EOM labelling. Arrowhead indicates an increased number of reporter positive cells surrounding the optic nerve (asterisk) at E12. (F-G’) Segmented periocular region. pom-ring (dotted lines) (n=3). **(H-H’)** Coronal sections at different eye levels of E11.75 *Tg:RARE^CreERT2^::R26^mTmG^* embryos immunostained for GFP (RA-responsive cells) and MYOD/MYOG (muscle). Higher magnification views as insets. Arrowheads mark presence of RA-responsive cells in medial periocular mesenchyme (H) and lateral-POM (H’). Asterisks indicate sporadic labelling in myogenic cells (n=3). **(I-J’)** EOM 3D-reconstructions of whole mount MF20 immunostaining of E12.5 of control (*Wnt1Cre: Rarb^fl/+^:Rarg^fl/+^*) (I,I’) and mutant (*Wnt1^Cre^:Rarb^fl/fl^:Rarg^fl/fl^*) (J,J’) embryos. Medial and lateral views are shown. Note absence of splitting in mutant EOM. Arrowheads indicate split EOM in controls (n=2 per genotype).

Compound *Aldh1a1/Aldh1a3* mutants (Matt et al., 2005; Molotkov et al., 2006), *Rarb/Rarg* mutants (Ghyselinck et al., 1997) and *Rarb/Rarg* NCC-specific mutants (Matt et al., 2008) display similar phenotypes in the anterior segment of the eye and POM. These findings have been attributed to the fact that ATRA synthesized by ALDH1A1 and ALDH1A3 in the neural retina activate RARB and RARG in the POM, which work as heterodimers with RXRA (Kastner et al., 1994; Mori et al., 2001). To directly examine whether the role of retinoic acid signaling in muscle patterning is driven non-cell autonomously, through the action of RARB/RARG receptors in the NCC-derived POM, we performed WMIF for muscle in *Wnt1^Cre^;Rarb^fl/fl^;Rarg^fl/fl^* embryos. Accordingly, these NCC-*Rarb/Rarg* mutants showed absence of muscle splitting (Fig 5I-J’) as observed in *Rdh10^-/-^* embryos (Suppl Fig 4F). Altogether, these data suggest that synthesis of ATRA from neural derivatives (retina, optic nerve, RPE) at early stages (≈E10.5) is crucial for EOM patterning, through its action on NCC-derived periocular connective tissues.

### Defective organization of the EOM insertions in retinoic acid signaling deficient embryos

To examine which connective tissue subpopulations, including the EOM insertions in the POM, were affected by retinoic acid signaling deficiency, we assessed the distribution of POM markers on tissue sections of *Aldh1a3^-/-^* and BMS493-treated embryos. Examination of the distrubution of Collagen XII (Fig 6A-C’), a marker of the sclera and the presumptive corneal epithelium (Oh et al., 1993), showed that ATRA-deficiency led to reduced expression of Collagen XII at E12 at the level of the medial-POM which gives rise to the sclera (Fig 6B-C, brackets, B’-C’ insets). In addition, while PITX2 was expressed continuously in the entire POM in controls (Fig 6D, dotted lines, D’, inset), expression in the medial-POM was lost upon ATRA-deficiency (Fig 6E-F, brackets, E’,F’, insets). This observation is in agreement with *Pitx2* being a retinoic acid signaling target in the NCC-derived POM (Matt et al., 2005; Matt et al., 2008; Molotkov et al., 2006) and the sclera being absent at fetal stages in *Rarb/Rarg*-NCC mutants (Matt et al., 2008). Interestingly, POM organization was unaffected in *Myf5^nlacZ/nlacZ^* mutants (Fig 6G, dotted lines, G’, inset), where the initial EOM anlage forms but myogenesis is aborted from E11.5 (Sambasivan et al., 2009). Thus, we conclude that intrinsic organization of the POM is dependent on retinoic acid signaling but not on the presence of myogenic cells.

**Fig 6.**
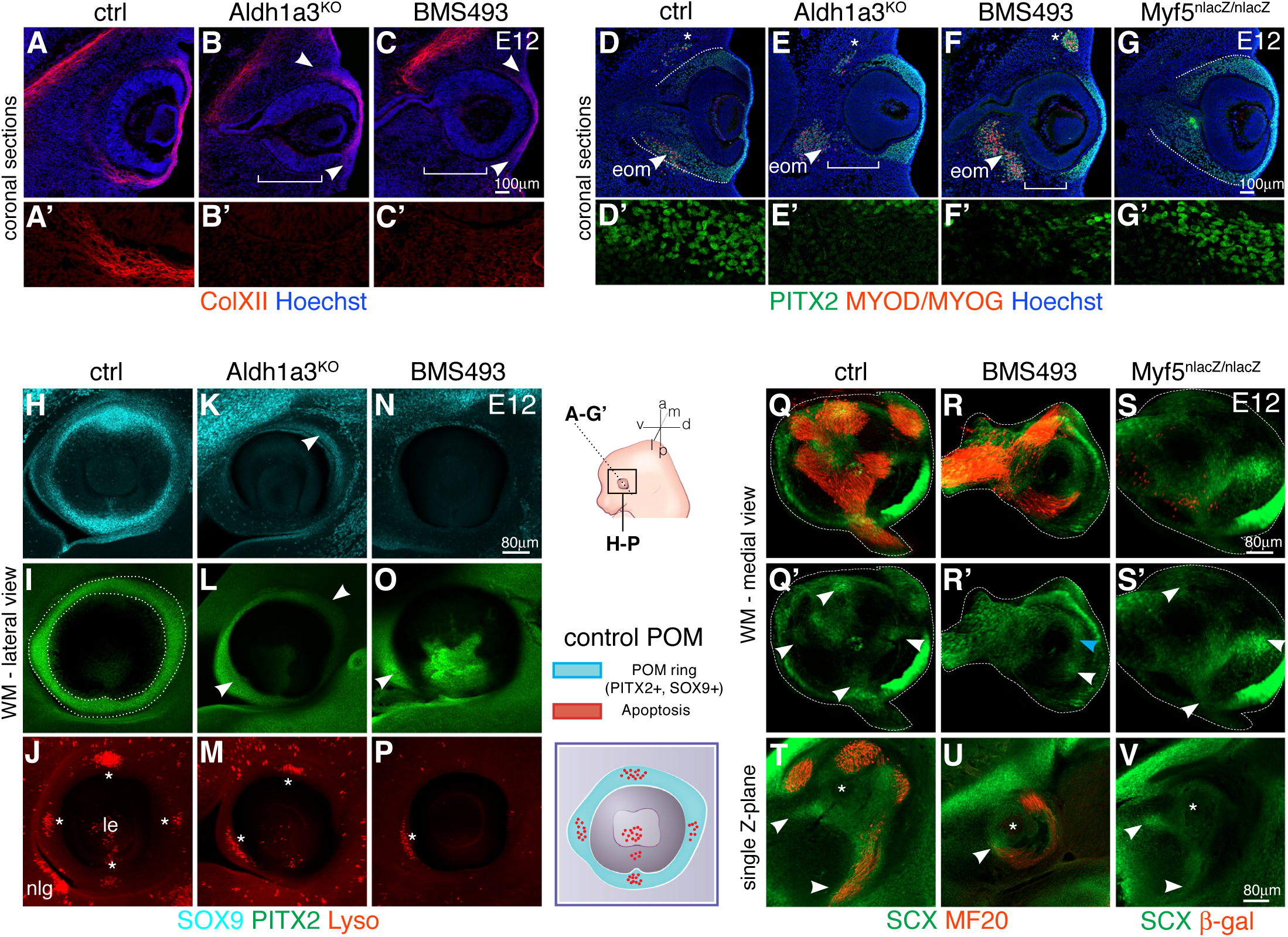
EOM insertions in the POM altered upon ATRA-deficiency. **(A-C’)** Immunostaining for Collagen XII (ColXII) on E12 coronal sections. Reduced ColXII expression domain in *Aldh1a3^KO^* (B,B’) and BMS493 treated embryos (C,C’) (arrowheads). ColXII expression in medial-pom (future sclera) is lost (brackets). Split ColXII channel as insets in (A’-C’). **(D-G)** Immunostaining for PITX2 (muscle progenitors, periocular mesenchyme) and MYOD/MYOG (myogenic cells) on coronal E12 sections. Asterisks indicate superior oblique muscle (SO). Dashed line mark periocular mesenchyme (pom). Brackets on *Aldh1a3^KO^* and BMS493 embryos indicate absence of PITX2 expression in medial-pom. Split PITX2 channel as insets in (D’-G’). **(H-P)** Lateral views of periocular region of whole mount immunostaining (GFP, SOX9, PITX2) and Lysotracker Red Staining (Lyso, apoptosis) of E12 control (H-J), *Aldh1a3^KO^* (K-M) and BMS493 treated (N-P) embryos (left eyes). *Aldh1a3^KO^* and BMS493 embryos do not show a complete pom-ring as controls (dotted lines in I). Arrowheads mark remaining expression of SOX9/PITX2 in the POM of *Aldh1a3^KO^* (L) or inhibitor treated (O) embryos. Asterisks in (J,M,P) mark apoptosis spots in POM. **(Q-V)** WMIF for MF20 (myofibers) and GFP (*Tg:Scx-GFP*) on control (Q,Q’,T), BMS493 treated (R,R’,U) and *Myf5^nlacZ/nlacZ^* E12 embryos (S,S’,V) (left eyes). **(Q-S)** The periocular area was segmented from adjacent structures for ease on visualization (dashed lines). **(T-V)** Single Z-section of the segmented volume. (n=3 per condition) White arrowheads in Q’,S’, T,V indicate correct position of tendon condensations for recti muscles. Blue arrowhead in R’ mark a diffuse *Scx-GFP*+ pattern at the position of the mispatterned MR (medial recti) muscle and white arrowheads in R’, U mark an ectopic *Scx-GFP* condensation at the muscle tip. le, lens; nlg, nasolacrimal gland; eom, extraocular muscles

To understand if the abnormal distribution of POM markers translates into EOM insertion defects, we examined the organization of the POM-ring in 3D in control, *Aldh1a3^-/-^* and BMS493-treated embryos. We systematically compared E11.75 and E12 embryos to visualize the organization of these tissues prior to, and during muscle splitting (Fig 2A-C). Control embryos showed the presence of partially overlapping SOX9 and PITX2 POM-ring domains, together with 4 apoptotic foci labelling the prospective tendon attachment points of the 4 recti muscles (Fig 6H-J, Suppl Fig 6A-C’,E,F). As expected, ATRA-deficiency perturbed these patterns in a dose dependent manner. In lateral views of the head of *Aldh1a3^-/-^* embryos, SOX9 and PITX2 had narrower expression domains than controls, restricted to the anterior-dorsal and ventro-lateral eye quadrants (Fig 6K-L, Suppl Fig 6G,H). In BMS493-treated embryos, expression of SOX9 in the POM (Fig 6N, Suppl Fig 6I) and RPE (Suppl Fig 6D,D’) was reduced to a minority of cells, and PITX2 expression was restricted ventro-laterally (Fig 6O, Suppl Fig 6J). Accordingly, the apoptotic foci were lost or disorganized in both cases of ATRA-deficiency with a single ventro-medial Lysotracker+ spot almost invariably present in both conditions (Fig 6M,P, Suppl Fig 6G-J). Finally, in medial views of the reconstructed 3D volume, we saw a continuum of PITX2 expression from the POM-ring to the medial-POM in control embryos (Suppl Fig 6N), while we noticed loss of medial-POM PITX2 expression in both cases of ATRA-deficiency (Suppl Fig 6O-P, blue arrowheads) as observed on tissue sections (Fig 6E’,F’, brackets). Thus, ATRA is required for the proper specification/organization of the medial POM and POM-ring.

As tendons were present at muscle tips in E13.5 *Aldh1a3^-/-^* and BMS493-treated embryos, but located ectopically with respect to a wild-type configuration (Fig 4F’-J’), we suspected that the initial 3D-organization of *Scx-GFP+* tendon and connective tissue progenitors might also be affected. We performed WMIF for differentiated muscle (MyHC) and tendon (*Tg:SCX-GFP*) and segmented the reconstructed 3D volume. In control embryos at E12.5, *Scx-GFP+* condensations projected from the tips of the future 4 recti muscles towards the POM-ring (Fig 6 Q,Q’, Video 7), facing the 4 apoptotic foci (Fig 6J, Video 3). Moreover, *Scx-GFP+* cells were also present in POM-muscle connective tissue presaging the final locales of the recti muscles (Fig 6T). In BMS493-treated embryos at E12, *Scx-GFP* expression was less organized and diffuse (Fig 6R,R’, blue arrowhead) yet *Scx-GFP*+ condensations could be observed at the tips of ectopic muscle masses (Fig 6R’,U white arrowhead, Video 8). In contrast, *Scx-GFP* prepatterning in muscle connective tissues was preserved in *Myf5^nlacZ/nlacZ^* mutants and *Scx-GFP+* condensations formed at the positions of prospective (although absent) recti muscles (Fig 6S,S’,V, white arrowheads). This situation is similar to the presence of individual tendons on the distal autopod of muscle-less limbs (Huang et al., 2015; Kardon, 1998) and branchial arches (Grenier et al., 2009). However, at later stages, tendons did not continue their maturation in the absence of EOMs (Suppl Fig 6Q-R’, video 9 and 10), reinforcing the notion that, as in most other anatomical locations (Huang, 2017), tendon differentiation and maintenance appear to ultimately rely on muscle-tendon interactions. In summary, ATRA produced by the developping eye influences the organization of the entire neural crest derived POM, including tendon condensations and EOM insertions in the POM-ring. Moreover, given that retinoic acid signaling exerts this action with a time interval that precedes and overlaps with that of EOM anlage segregation into individual muscle masses, we propose that EOM and insertion site patterning are regulated in a coordinated manner.

## DISCUSSION

The stark differences in embryological origins, function and susceptibility to disease among cranial and somite-derived muscles (Sambasivan et al., 2011), provides impetus to study in detail the morphogenesis of different muscle groups and their integration as part of the musculoskeletal system. Here we focused on EOMs, an evolutionarly conserved group of muscles that are precisely engineered for fine displacement of the eyeball, and thus crucial for visual acuity. Using genetic and 3D imaging approaches, we analysed EOM development from their emergence as a unique anlage to the establishment of a fully formed functional unit with insertions in the sclera, which sets it apart from classical muscle-to-bone insertions studied to date. We identified a spatiotemporal window in which retinoic acid signaling from the target organ is required for patterning of NCC-derived soft tissues of the POM. These findings provide insights into the deployment of site-specific programs for the establishment of anatomically distinct muscle functional units, with precise definition of muscle shapes and topographical wiring of their tendon attachments.

### Dual origin of EOM insertions

The connective tissues of the rostral cranium (bone, cartilage, tendon and muscle connective tissue) were reported to be derived from the neural crest (Nassari et al., 2017; Noden and Francis-West, 2006; Ziermann et al., 2018). However, the hypochiasmatic cartilages, from which the recti muscles originate in mammals, are an exception to this rule as they are mesoderm-derived (McBratney-Owen et al., 2008). Here, we found that at this location, tendon and muscle connective tissue are also of mesodermal origin. As such, EOM originate in mesoderm-derived bone and insert in NCC-derived fibrous tissue of the sclera with muscle connective tissue following the embryonic origins of the respective attachments as observed in other anatomical locations (Heude et al., 2018). Taken together, our observations redefine a novel boundary for NCC contribution to the connective tissues of the periocular region and suggest that at their origin, the EOMs and their associated connective tissues might have evolved neomorphically *in situ* from the same mesodermal source.

### RA responsiveness on the POM

Previous studies showed that NCC have a critical role in the acquisition of cranial muscle morphology (Bohnsack et al., 2011; McGurk et al., 2017; Shimizu et al., 2018; Tokita and Schneider, 2009; von Scheven et al., 2006). Our data using the *Rare^CreERT2^* reporter and following invalidation of *Rarb/Rarg* in NCC derivatives suggest that the action of retinoic acid signaling on EOM patterning is also indirect, through its action on periocular connective tissues.

The observation that BMS493 treatments to impair ATRA signaling results in a more severe phenotype than that observed in *Aldh1a3* mutants could have several explanations. First, BMS493 is not a RAR antagonist but pan-RAR inverse agonist, which is capable of repressing RAR basal activity by favoring the recruitment of corepressors and promoting chromatin compaction (Germain et al., 2009). Second, additional sources of ATRA could signal to the POM and be inhibited by the BMS493 treatment. Candidates include *Aldh1a1* (retina), *Aldh1a2* (temporal mesenchyme) and *Cyp1B1*, a member of the cytochrome p450 family of mono-oxygenases which can generate ATRA in the retina in a retinaldehyde dehydrogenease-independent manner (Chambers et al., 2007). Noneless, the severe patterning phenotypes observed in the *Aldh1a3* single mutant suggest that the local synthesis of ATRA in the RPE, optic stalk and ventral retina at early stages (≈E10.5) cannot be fully compensated by the action of other retinaldehyde dehydrogeneases in adjacent tissues (this study, (Dupe et al., 2003; Harding and Moosajee, 2019)). Interestingly, the constitutive *RARE-hsp68-lacZ* transgene displays strong reporter activity in the retina and RPE (Matt et al., 2005; Rossant et al., 1991). Thus, we cannot exclude the possibility that some of the phenotypes observed in periocular connective tissues of the *Aldh1a3* mutant are partly due to changes in signalling pathways downstream of ATRA in the retina and/or RPE, that subsequently act in the POM.

### EOM and tendon patterning are concerted processes under RA control

We identified a short temporal window (E10.5-E11.75) where ATRA is essential for the precise morphogenesis of several elements of the POM, including the formation of tendon condensations and insertion sites in the POM-ring. These phenotypes are already evident from E11.75, a stage where the EOM still figures as a muscle anlage. Interestingly, analysis of chick and mouse limb development also identified a short time window where limb mesenchyme perturbations result later in muscle and tendon patterning defects (Chevallier and Kieny, 1982; Hasson et al., 2010). Moreover, concomitant muscle and tendon defects have been observed in the chick limb and zebrafish jaw upon retinoic acid signaling misregulation (McGurk et al., 2017; Rodriguez-Guzman et al., 2007). Altogether, these results suggest that at certain anatomical locations including the EOMs, endogenous local variations in the concentration of retinoids contribute to the establishment of *Scx-GFP+* condensations, a critical step in muscle functional unit assembly. As ectopic muscles masses are associated with ectopic tendons upon ATRA deficiency, one possible explanation is that retinoic acid signaling directly controls *Scx-GFP+* condensations, which in turn help organize the adjacent forming muscles. On the other hand, it is also possible that connective tissue mispatterning upon ATRA deficiency determines the organization of the muscle masses and the muscle tips direct the organization of the *Scx-GFP+* condensations.

### Role of RA in the integration of EOM patterning and insertion site formation

The EOMs insert into the sclera, a non-bone structure in mammals. The generation of 4 attachment points that precisely mirror the position of the 4 recti muscles is a morphogenesis conundrum, as details on their specification are scarse. Four mesenchymal condensations have been described at the periphery of the developing optic cup in cat and human embryos, in positions facing the locations of the recti muscles (Gilbert, 1957; Gilbert, 1947). In the mouse, at equivalent developmental timepoints (E11.5-E12.5) and homologous positions, apoptotic foci were observed and suggested to provide attachment for the recti muscles (Sulik et al., 2001). Using 3D-imaging, we showed that at E11.75, the apoptotic foci are aligned with *Scx-GFP*+ condensations that project medially, before any sign of splitting of the EOM anlage. As the apoptotic foci decrease in size from E12.5 onwards concomittant with ongoing muscle splitting and tendon formation, it is possible that they initially presage the POM for future tendon insertions, but are not required for attachment *per se*. These foci appear in a SOX9+ PITX2+ domain of the future sclera, however, to our knowledge their presence has not been reported in superstructures at other anatomical locations.

In agreement with the role of RA in the induction of cell death in another locations in the embryo (Dupe et al., 2003), our study shows that upon perturbation of RA signaling, the apoptotic foci in the POM are absent or severely reduced. However, it is also possible that the loss of foci upon RA inhibition is secondary to loss or mispatterning of the SOX9+ PITX2+ POM-ring. Thus, it will be of interest to determine if null or NCC specific mutations in PITX2 affect the patterns or amount of naturally occurring cell death in the periocular condensations. Although our work sheds light on the genetic mechanisms that regulate EOM insertion, the cellular mechanisms underlying this process remains undefined. It is tempting to speculate that the apoptotic foci are related to the disappearance of signaling centers (Jernvall et al., 1998; Nonomura et al., 2013) or compaction-mediated cell death and extrusion (Moreno et al., 2019). Future experiments directed towards modifying the amount or timing of cell death will be informative.

Few reports have demonstrated that mispatterning of specific muscles is coupled with aberrant superstructures (Colasanto et al., 2016; Swinehart et al., 2013; Tokita and Schneider, 2009) or removal of the prospective tendon attachment sites (Rodriguez-Guzman et al., 2007). Our results, are in agreement with that model given that EOM mispatterning is concomitant with aberrant EOM insertions. Given that mammals do not develop a cartilage layer within the sclera (Franz-Odendaal and Vickaryous, 2006; Thompson et al., 2010), expression of SOX9 in the POM-ring of mouse embryos at the time of EOM patterning is intriguing. SOX9 expression at this level appears to be dependent on a functional RPE and ATRA ((Thompson et al., 2010), this study). In this context, our data suggest that transient SOX9 expression may represent a redeployment of the developmental module for tendon attachment, despite the fact there is no cartilage in the mammalian sclera. Finally, transcriptome analysis revealed the existence of global and regional regulatory modules for superstructure patterning in the limb bones (Eyal et al., 2019), offering a mechanism to induce variations in attachment sites without having to rewrite the entire skeletogenic program (Eyal et al., 2019). As several markers of the POM (*Pitx2*, *Foxc1/2*) and retinoic acid signaling modulators (*Cyp26a1/b1*) were identified as part of specific limb superstructure signatures (Eyal et al., 2019), it is tempting to speculate that those genes also play a conserved role in the generation of the attachment module of the EOMs.

## Conclusion

The developing eye has been proposed to be key organizer of craniofacial development, independent of its role in vision (Kish et al., 2011). This notion is based on the role of the eye for proper NCC migration to the periocular region (Langenberg et al., 2008), the common association of ocular and craniofacial developmental abnormalities (Zein, 2012) and the initial development of an eye in blind vertebrate species (Durand, 1976; Kish et al., 2011; Yamamoto et al., 2004). Moreover, as the appearence of a “camera-type” eye is a vertebrate innovation (Lamb et al., 2007) and the EOM are already present in lamprey (Suzuki et al., 2016), it is possible that muscles and their target tissue might have co-evolved. Here, by characterizing coordinated patterning of the EOMs, their respective tendons and attachments, our findings illustrate further the role of the developing eye as a signaling center allowing integration of the EOM functional unit in the POM. Our results show that the tissue interactions during the development of this craniofacial muscle unit share features with those described in the limb, but with additional regional properties (e.g. cell death, specification of attachments in the sclera) that seem to have been specifically incorporated into this group. Moreover, the capacity to instruct muscle patterning through variations in connective tissue derivatives, provides a mechanism to explain the plasticity of the musculoskeletal system at the anatomical and inter-species levels, while ensuring structural and functional integration during evolution. These findings also imply that the generation of musculoskeletal units do not require major restructuring of the developmental programs of all the tissues implicated. Instead, co-option of a general program and simultaneous addition of local features appear to elaborate musculoskeletal diversity.

## Supplementary Figure Legends

**Supplementary Fig 1.**
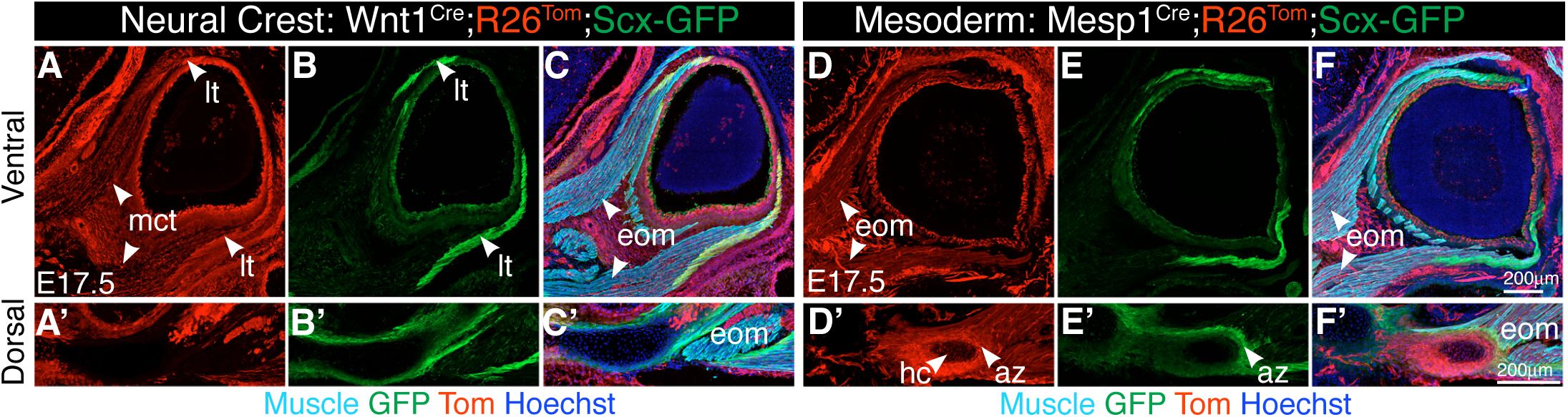
Lineage contributions to the EOM functional unit. (A-F) Neural Crest cell (NCC, *Wnt1^Cre^;R26^Tom^*) and mesoderm (*Mesp1^Cre^;R26^Tom^*) lineage contributions to the periocular region on coronal cryosections of E17.5 embryos, combined with immunostaining for GFP (*Tg:Scx-GFP* tendon reporter) and muscle (Tnnt3). Sections at ventral (A-F) and dorsal (A’-F’) levels. Note that lateral tendons at the level of the orbit are NCC-derived (A-C), whereas tendons at the levels of the annulus of zinn (D’-F’) are mesoderm-derived. az, annulus of zinn; eom, extraocular muscles; hc, hypochiasmatic cartilage; lt, lateral tendon (at the level of the insertion); mct, muscle connective tissue

**Supplementary Fig 2.**
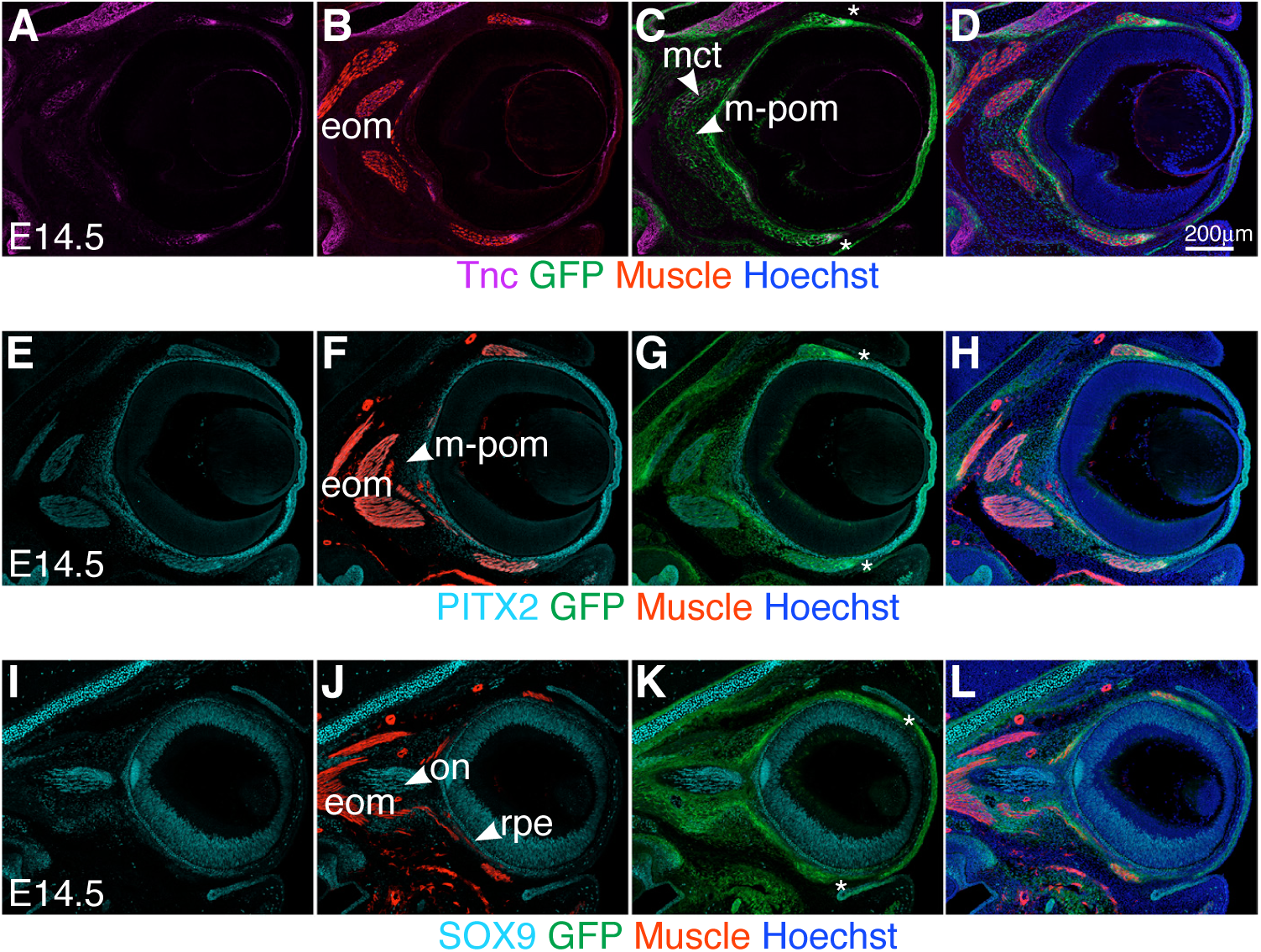
Developmental timing of EOM functional unit components. (A-L) Immunostaining for the indicated markers on coronal sections of E14.5 *Tg:Scx-GFP* embryos. Tnc (tenascin) (n=2 per condition). eom, extraocular muscles; lt, lateral tendon (at the level of the insertion; asteriks); m-pom, medial periocular mesenchyme; mct, muscle connective tissue; on, optic nerve; rpe, retinal pigmented epithelium

**Supplementary Fig 3.**
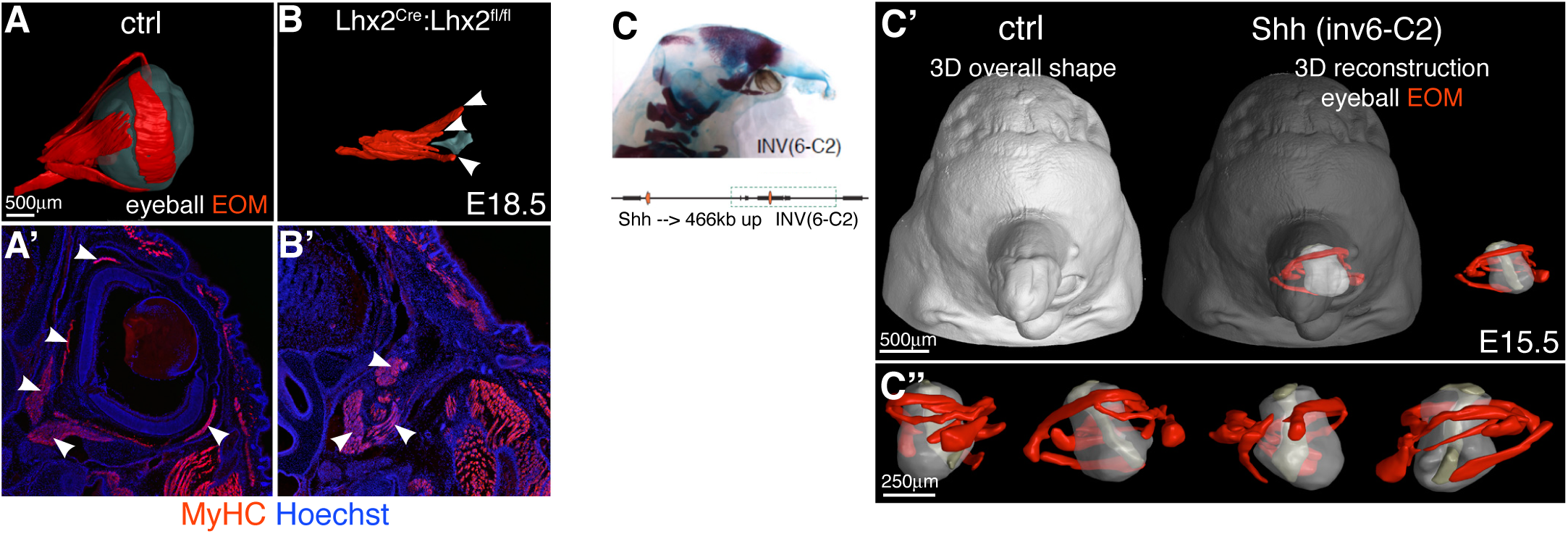
EOM morphogenesis in mutants with eye defects. (A-B) µCT-based 3D-reconstruction of EOM and eyeball in E18.5 control (A) and *Lhx2Cre:Lhx2fl/fl* (B) embryos. Arrowheads highlight some extent of EOM segregation in the mutant. (A’-B’) Coronal sections of control (A’) and mutant (B’) embryos stained with MyHC. Arrowheads indicate individual EOM masses (n=2). **(C-C’’)** Analysis of EOM patterning in E15.5 embryos containing inversions of *Shh* genomic regulatory regions (Inv(6-C2)). (C) Skeletal preparation of mutant embryos displaying cyclopia. (C’-C’’) µCT-based 3D reconstruction of EOM and eyeball of mutant embryos (n=2).

**Supplementary Fig 4.**
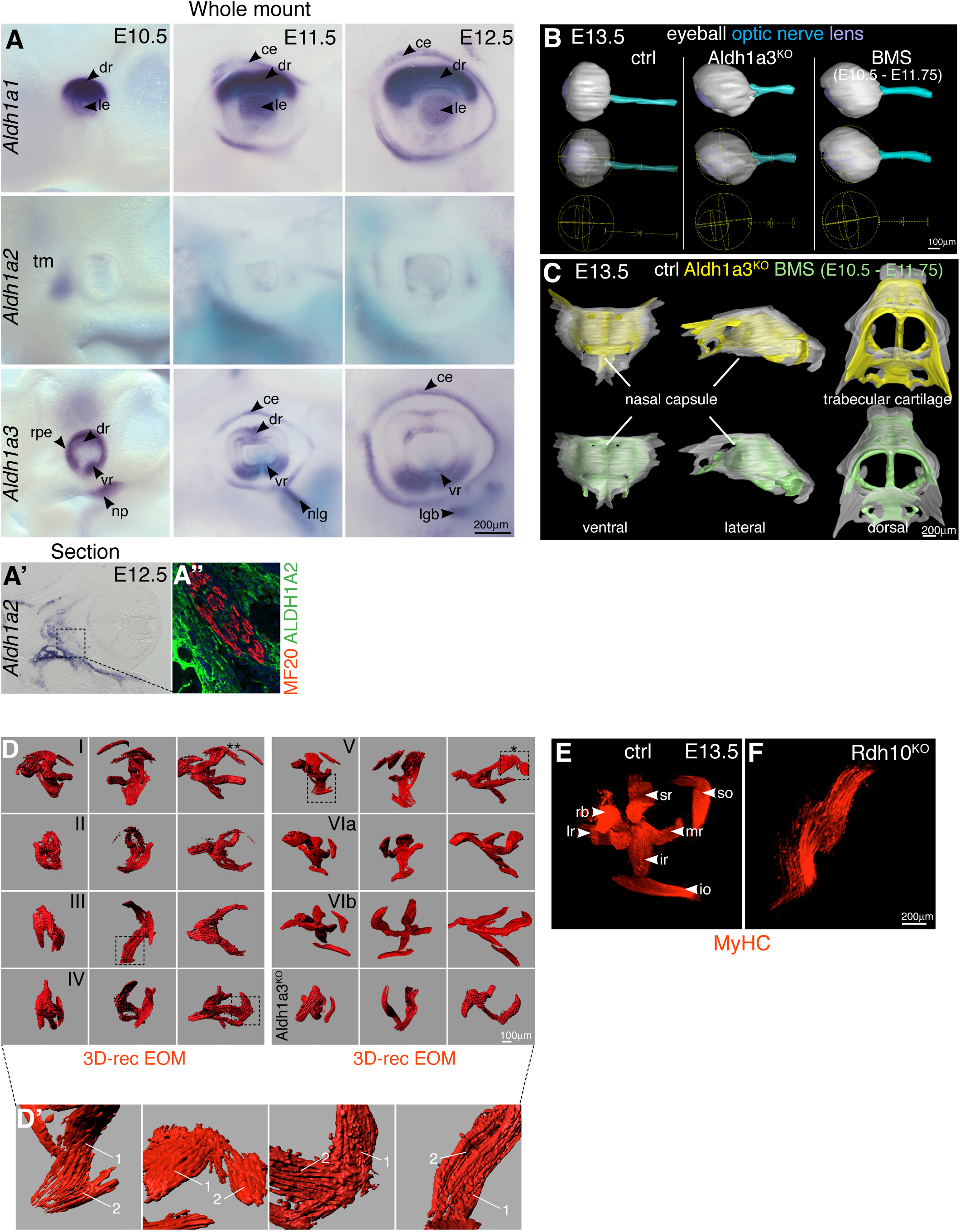
EOM morphogenesis is dependent retinoic acid signaling. (A)Whole mount in situ hybridization for *Aldh1a1*, *Aldh1a2* and *Aldh1a3* in E10.5, E11.5 and E12.5 embryos (n=3). **(A’)** In situ hybridization for *Aldh1a2* and immunostaining for ALDH1A2 and muscle (MF20) on E12.5 coronal sections. ALDH1A2 is expressed in the temporal mesenchyme and muscle connective tissue (n=3). **(B)** µCT-based 3D-reconstruction of eyeball, optic nerve and lens of E13.5 control, *Aldh1a3^KO^* and BMS treated embryos (n=2 each genotype). The lower row is a scheme of a sphere fitting the eyeball and lens, and a cylinder for the optic nerve. Note ventralization of the eyeball in *Aldh1a3^KO^* and BMS treated embryos. **(C)** µCT-based 3D-reconstruction of the mesenchymal condensations of the nasal capsule and trabecular cartilage of E13.5 control (white), *Aldh1a3^KO^* (yellow) and BMS-treated (green) embryos. **(D-D’)** EOM 3D-reconstructions of whole mount MyHC immunostaining of E13.5 of BMS-treated embryos as described in Table S1. The most severe phenotype obtained in each condition is shown. Two different examples of the phenotype observed upon treatment VI (a,b) are shown (most and least severe). An *Aldh1a3^KO^* embryo is shown as comparison. Asterisk denote ectopic duplicated SO (superior oblique) muscle. (D’) Higher magnification views from D with examples of adjacent, non-split muscle masses (1,2) depicted by differential fiber orientation. **(E-F)** Whole mount MyHC immunostaining of E13.5 control (G) and *Rdh10^KO^* (H) embryos. Note total absence of muscle splitting in mutant (n=3 per condition). ce, corneal ectoderm; dr, dorsal retina; io, inferior oblique; ir, inferior rectus; le, lens; lgb, lacrimal gland bud; lr, lateral rectus; mr, medial rectus; nlg, nasolacrimal groove; np, nasal pit; os, optic stalk; se, surface ectoderm; so, superior oblique; sr, superior rectus; rb, retractor bulbi; rpe, retinal pigmented epithelium; tm, temporal mesenchyme; vr, ventral retina

**Supplementary Fig 5.**
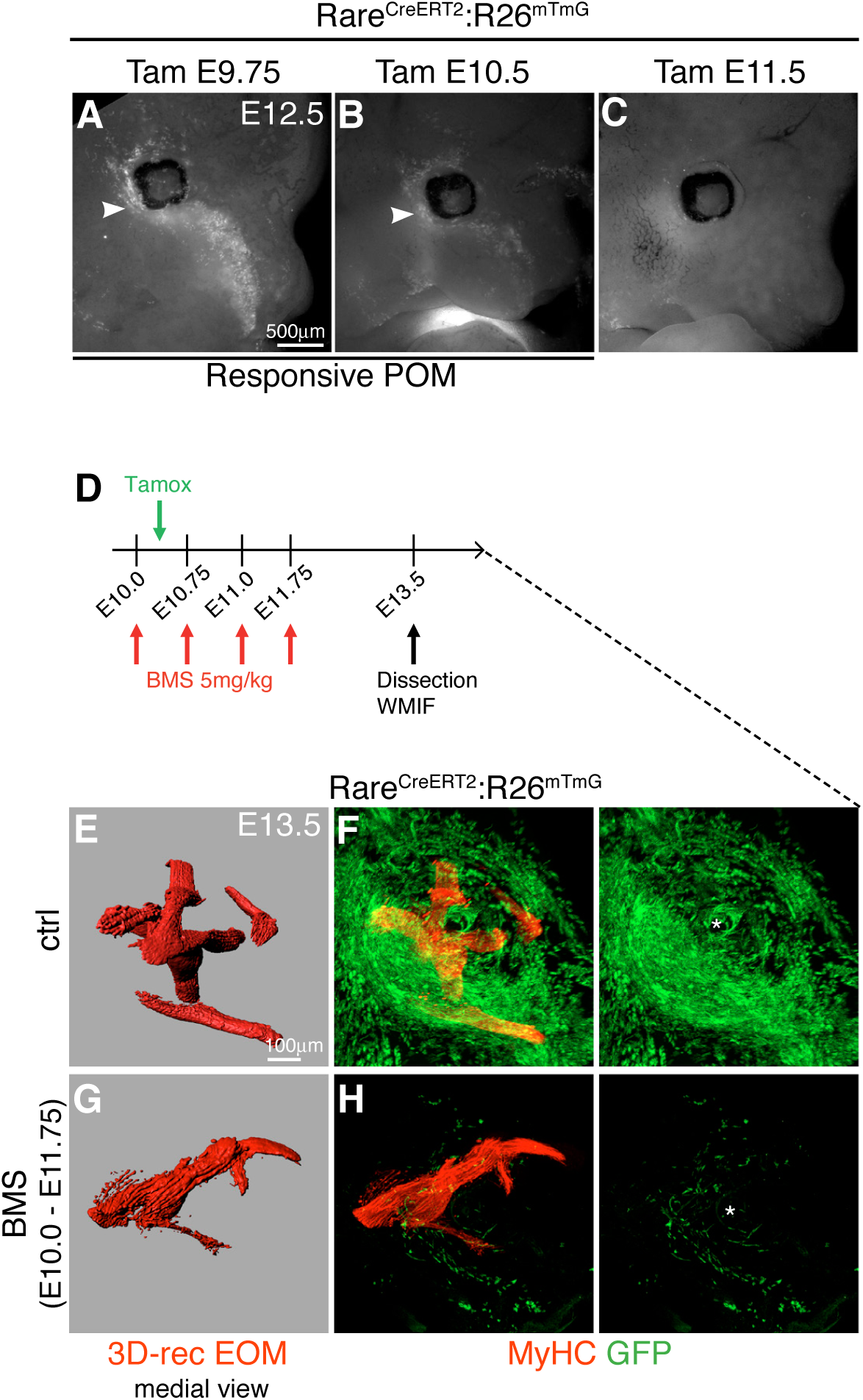
Retinoic acid signaling responsiveness in the periocular region. **(A-C)** Macroscopic views of endogenous GFP fluorescence of *Tg:RARE^CreERT2^:R^26mTmG^* embryos. Tamoxifen (Tam) was injected into pregnant females at indicated time points. Arroheads indicate labelling in periocular mesenchyme (POM) (n>3 per condition). **(D)** Strategy used to determine responsiveness of *Tg:RARE^CreERT2^* reporter in presence of RA signaling reverse agonist BMS493. BMS493 was injected to pregnant females between E10 and E11.75. **(E-H)** Whole mount immunostaining for MyHC (muscle), GFP (ATRA-responsive cells) of control (E, F) and BMS493-treated embryos (G, H). (E, G) Medial views of 3D-reconstructed EOM labelling. BMS493 treatment before and after tamoxifen induction reveals a drastic decrease in GFP+ cells in the periocular region (H) compared to controls (F). Asterisks mark the location of the optic nerve (n=2).

**Supplementary Fig 6.**
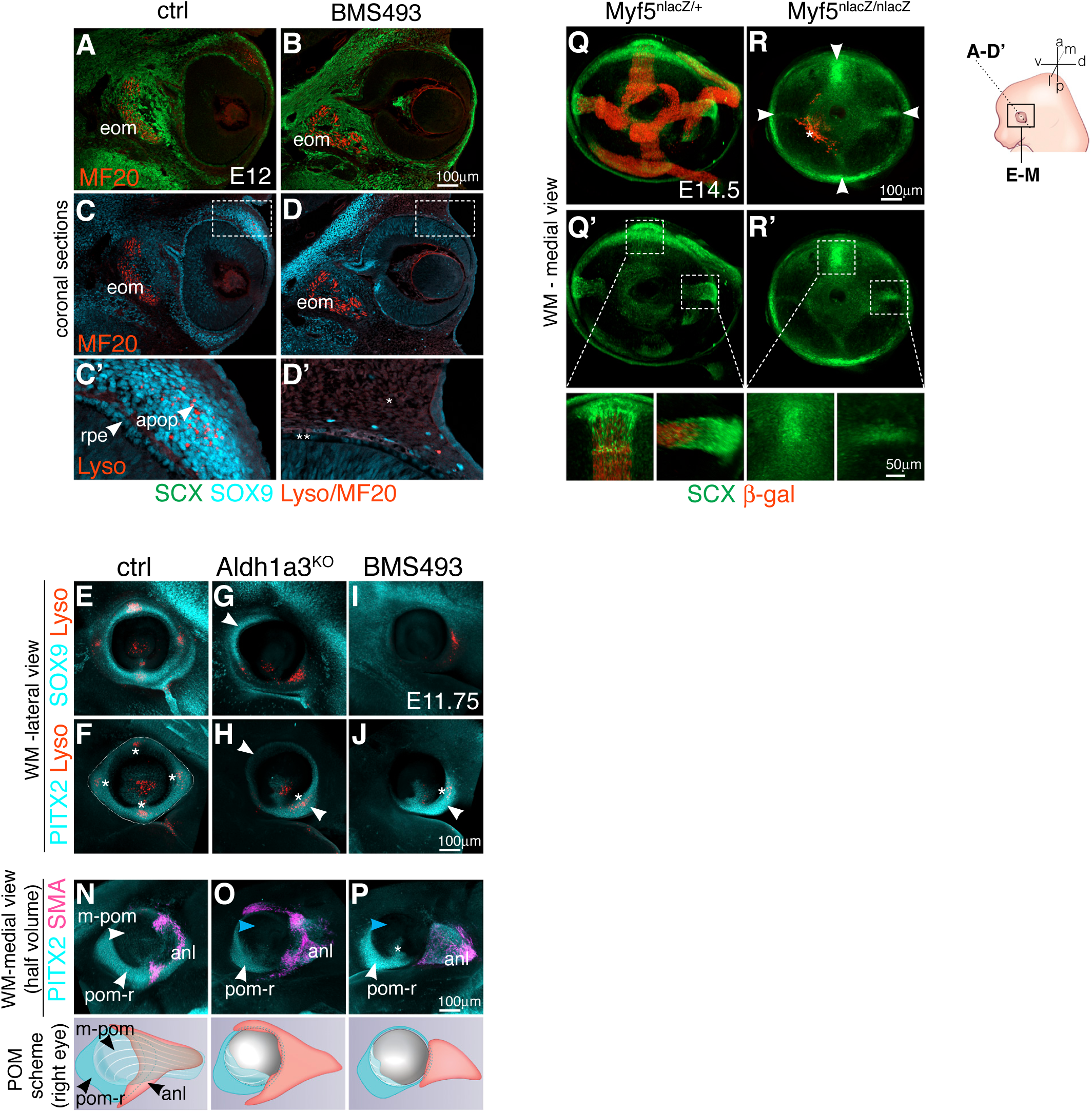
EOM insertions in the POM are altered upon ATRA-deficiency. **(A-D)** Immunostaining for GFP (tendon progenitors), SOX9 (periocular mesenchyme, POM), MF20 (myofibers) together with Lysotracker Red staining (Lyso, apoptosis) on coronal E12 sections of *Tg:Scx-GFP* control (A-B’) and BMS493 treated (C-D’) embryos. (B’,D’) higher magnification views of the periocular mesenchyme (POM) region. In BMS493-treated embryos, Lysotracker Red and SOX9 staining are absent in POM (D’, asterisk) and SOX9 staining also missing in the RPE (D’, double asterisk). **(E-P)** Whole mount immunostaining for SOX9 and PITX2, SMA and Lysotracker Red Staining of E11.75 control (E, F, N), *Aldh1a3^KO^* (G,H, O) and BMS493-treated (I, J, P) embryos (right eyes). In lateral views, neither *Aldh1a3^KO^* or BMS493 embryos show a PITX2+ pom-ring as the controls (F). Arrowheads in (G,H,J) mark remaining expression of SOX9/PITX2 in POM of mutant or inhibitor treated embryos. Asterisks in (F,H,J) mark apoptosis spots in POM. **(N-P)** Segmented medial views of periocular region of control embryos. Volumes were truncated in Z for clarity. Full view as schemes below. The PITX2+ pom-ring is continuous with the medial-pom (white arrowhead, N). In mutant and inhibitor treated embryos, residual PITX2 expression in POM is discontinous with the medial-POM. **(Q-R)** Whole mount immunostaining for GFP and β-gal (*Myf5^nlacZ^* reporter, myogenic progenitors) of E14.5 *Tg:Scx-GFP*::*Myf5^nlacZ/+^* (Q, control) and *Tg:Scx-GFP*::*My5^nlacZ/nlacZ^* (R, mutant) embryos (left eyes). Asterisk indicate few remaining β-gal+ cells in mutant. (Q’,R’) split GFP channel. Arrowheads in R indicate the correct position of tendon condensations for the 4 recti muscles although these are absent in the mutant. Lower panel, higher magnification views. m, extraocular muscle; anl, anlage; apop, apoptosis; m-pom, medial periocular mesenchyme; l-pom, lateral periocular mesenchyme; rpe, retinal pigmented epithelium.

### Videos

**Video 1. Temporal sequence of EOM patterning.** 3D-recontruction of whole mount immunostaining of the developing EOM in embryos ranging from E11.5 to E14.5. EOM were segmented from adjacent head structures and 3D-reconstructed in Imaris.

**Video 2.** 3D-recontruction of EOM functional unit at E11.75. Immunostaining for Muscle (MF20, Cyan) and tendon (*Tg:Scx-GFP*, Green). Apoptotic foci are visualised by Lysotracker Red. A clipping plane is added for clarity of visualisation. Note how tendon condensations start to organize radially and towards the apoptotic foci in the POM.

**Video 3.** 3D-recontruction of EOM functional unit at E12.5. Immunostaining for Muscle (MF20, Cyan) and tendon (*Tg:Scx-GFP*, Green). Apoptotic foci are visualised by Lysotracker Red. A clipping plane is added for clarity of visualisation. Note how tendon condensations of the recti muscles are more refined at this stage and project towards the apoptotic foci in the POM.

**Video 4.** 3D-recontruction of te EOM functional unit at E13.5 of a BMS treated embryo (E10.5→ E11.75). Immunostaining for Muscle (MyHC, Red) and tendon (*Tg:Scx-GFP*, Green). Note the presence of tendon condensations at the tips of mispatterned muscle masses.

**Video 5.** 3D-recontruction of whole mount immunostaining of control EOM (MyHC, Red) and *Tg:Rare^CreERT2^:R26^mTmG^* reporter (GFP, Green) at E13.5 (Tamoxifen induction at E10.5). EOMs are shown as isosurface for clarity of visualisation.

**Video 6.** 3D-recontruction of whole mount immunostaining of EOMs (MyHC, Red) and *Tg:Rare^CreERT2^:R26^mTmG^* reporter (GFP, Green) at E13.5 (Tamoxifen induction at E10.5) on BMS-treated embryos (E10.5 → E11.75). EOMs are shown as isosurface for clarity of visualisation.

**Video 7.** 3D-recontruction of whole mount immunostaining of control EOM (MyHC, Red) and tendon (*Tg:Scx-GFP*, Green) at E12.5. A clipping plane has been added for clarity of visualisation.

**Video 8.** 3D-recontruction of whole mount immunostaining of EOM (MyHC, Red) and tendons (*Tg:Scx-GFP*, Green) of a BMS-treated embryo (E10.5 → E11.75) at E12.5. A clipping plane has been added for clarity of visualisation. Note that distribution of Scx+ cells is more difuse than for controls (Video 7), but an *Scx-GFP*+ condensation is present at the tip of the inferior muscle mass.

**Video 9.** 3D-recontruction of whole mount immunostaining of control EOM (MYOG/MYOG, Red) and tendon (*Tg:Scx-GFP*, Green) at E14.5.

**Video 10.** 3D-recontruction of whole mount immunostaining of *Myf5^nlacZ/nlacZ^* EOM (MYOG/MYOG, Red) and tendon (*Tg:Scx-GFP*, Green) at E14.5.

## Materials and methods

### Mouse strains and animal information

Animals were handled as per European Community guidelines and the ethics committee of the Institut Pasteur (CETEA) approved protocols. The following strains were previously described: *Mesp1^Cre^* (Saga et al., 1999), *Tg:Wnt1^Cre^* (Danielian et al., 1998), *Tg:Rare^CreERT2^* (Schedl et al., unpublished), *R26^Tom^* (Ai9; (Madisen et al., 2010)), *R26^mTmG^* (Muzumdar et al., 2007), *Tg:Scx-GFP* (Pryce et al., 2007), *Myf5^nlacZ^* (Tajbakhsh et al., 1996), *Aldh1a3^KO^* (Dupe et al., 2003), *Rarb^flox^* (Chapellier et al., 2002a), *Rarg^flox^*, (Chapellier et al., 2002b), *Rdh10^KO^* (Rhinn et al., 2011), *Pax6^Sey/Sey^* (Roberts, 1967). *Rdh10^KO^* embryos were received from the laboratory of Pascal Dollé. *Wnt1Cre:Rarb^fl/fl^:Rarg^fl/fl^* embryos were received from the laboratory of Valerie Dupé. *Pax6^Sey/Sey^* and control embryos were received from the laboratory of James Briscoe. *Lhx2^Cre^;Lhx2^fl/fl^* (Hagglund et al., 2011) were received from the laboratory of Leif Carlsson. *Shh(invC-6)2* embryos were received from the laboratory of François Spitz.

To generate experimental embryos for *Mesp1^Cre^* or *Wnt1^Cre^* together with *Tg:Scx-GFP* and *R26^Tom^* lineage tracings, Cre/+ males were crossed with *Tg:Scx-GFP::R26^Tom/Tom^* females. Mice were kept on a mixed genetic background C57BL/6JRj and DBA/2JRj (B6D2F1, Janvier Labs). Mouse embryos and foetuses were collected between embryonic day (E) E10 and E18.5, with noon on the day of the vaginal plug considered as E0.5. Pregnant females were euthanised by cervical dislocation.

To induce recombination with the *Rare^CreERT2^:R26^mTmG^* line, 5mg of tamoxifen (Sigma #T5648) were administered by gavage to pregnant females. A 25mg/ml stock solution in 5% ethanol and 95% sunflower seed oil was prepared by thorough resuspention with rocking at 4°C.

To inhibit retinoic acid signaling, pregnant females of relevant genotypes were injected intraperitoneally with 10mg/kg of BMS493 (Tocris, 3509), a pan-retinoic acid receptor (pan-RAR) inverse agonist. A 5mg/ml BMS493 stock solution in DMSO (SIGMA, D2650) was prepared and stored in single use aliquots at -20°C in tight cap tubes. At the time of injection, the aliquot was thawed, 200ul of sterile PBS was added per 50ul aliquot and injected immediately.

### Immunofluorescence

Embryos were fixed 2.5h in 4% paraformaldhehyde (PFA; 15710, Electron Microscopy Sciences) in PBS with 0,2-0,5% Triton X-100 (according to the embryonic stage) at 4°C and washed overnight at 4°C in PBS. For cryosectioning, embryos were equilibrated in 30% sucrose in PBS overnight at 4°C and embedded in OCT. Cryosections (16-18µm) were allowed to dry at RT for 30 min and washed in PBS. Immunostaining was performed as described in (Comai et al., 2019). TUNEL staining, that marks double strand breaks, was performed with the In Situ Cell Death Detection Kit/Fluorescein (Roche, 11 684 795 910). Slides were first pretreated with a 2:1 mix of Ethanol:Acetic Acid 5min -20C, washed twice 20 min with PBS at RT and processed for TUNEL staining as described by the manufacturer. For whole mount immunostaining, embryos were fixed and washed as above and dehydrated in 50% Methanol in PBS and twice in 100% Methanol, 30min each at RT and kept at -20°C till needed. Heads were rehydrated, the periocular region was micro-dissected in PBS and immunostaining performed as described in (Comai et al., 2019). For embryos older than E13.5, the alternative pretreatment (containing 0.1%Tween-20, 0.1%TritonX100, 0.1%Deoxycholate, 0.1%NP40, 20%DMSO in PBS) and primary antibody immunolabelling steps of the idisco protocol (https://idisco.info/idisco-protocol/) were generally used. In all cases secondary antibodies were applied in blocking buffer as described in (Comai et al., 2019) for >4 days at 4C with rocking. After immunolabelling, samples washed in 0.1%Tween/PBS, dehydrated in 50% Methanol in PBS, 100% Methanol 10min each at RT, cleared with a mix benzyl alcohol and benzyl benzoate (BABB) and mounted for imaging as described in (Yokomizo et al., 2012).

For detection of cell death in whole mount live tissues, LysoTracker Red staining was done, as it reveals lysosomal activity correlated with increased cell death (Fogel et al., 2012; Zucker et al., 1999). Embryos were quickly dissected in HBSS (Invitrogen, 14025-092), incubated in 2ml tubes containing 5µM of LysoTracker™ Red DND-99 (Molecular Probes, L7528) 45min at 37°C with rocking in the dark, washed twice in PBS, fixed and processed for cryosections or whole mount immunostaining as above.

### Antibodies

Primary and secondary antibodies used in this study are listed in Table 2. To detect differenting EOM we used: a-smooth muscle actin (SMA), which is transiently expressed in differentiating myoblasts and myotubes (Babai et al., 1990; Li et al., 1996; Sawtell and Lessard, 1989); Desmin, an early cytoskeletal muscle protein expressed in myoblasts, myotubes and myofibers (Babai et al., 1990; Li et al., 1993) and myosin heavy chain (MF20) to label sarcomeric myosin (Bader et al., 1982).

**Table 2.**
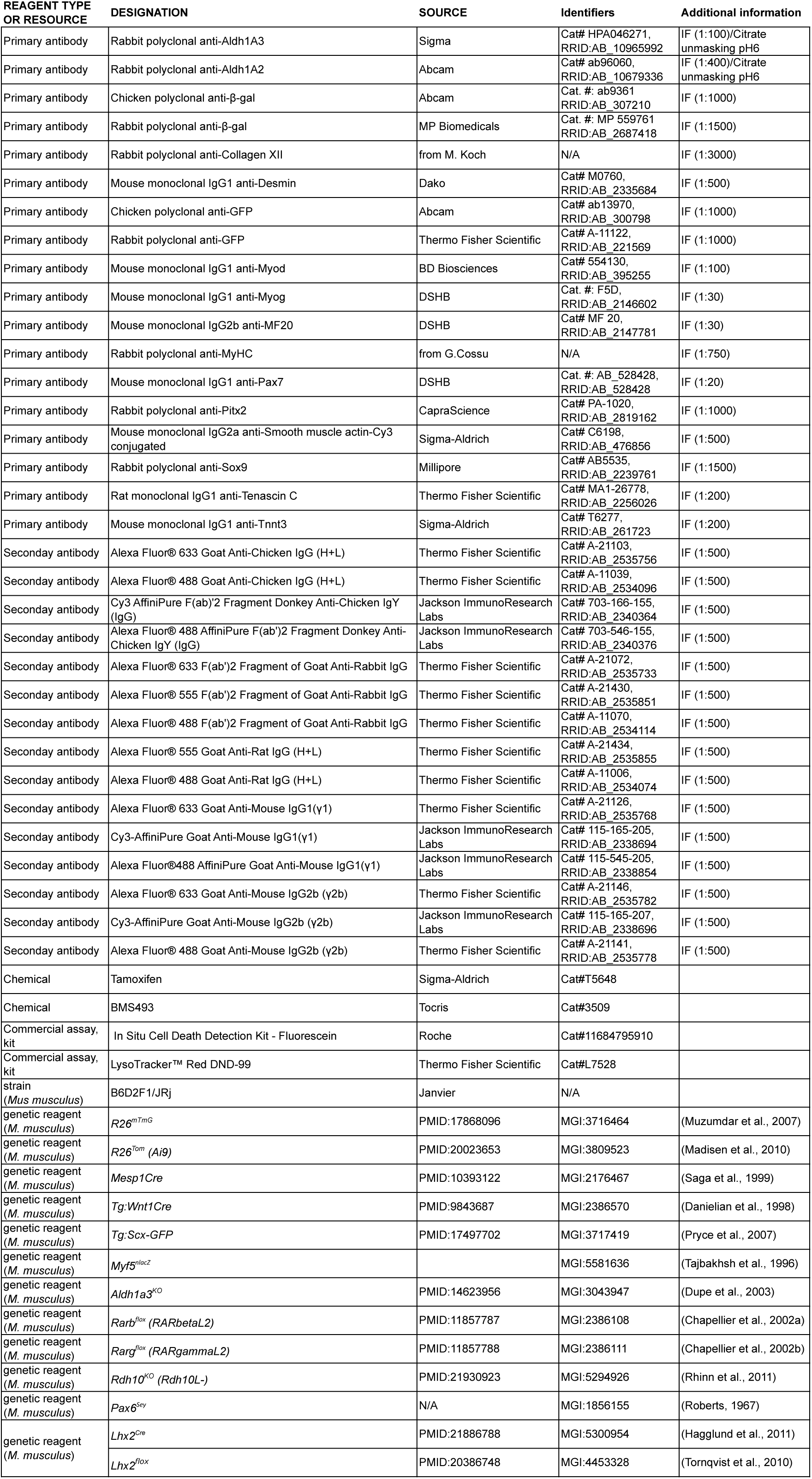
Antibodies and resources used in this study.

### Static Imaging

A Zeiss SteREO Discovery V20 macroscope was used for imaging the endogenous fluorescence of whole embryos at the time of dissection. For tissue sections and whole mount immunostaining of cleared embryos a LSM700 and a LSM800 laser-scanning confocal microscope with ZEN software (Carl Zeiss) were used.

All images were assembled in Adobe Photoshop and InDesign (Adobe Systems). Volume-3D rendering of the z-stack series was performed in Imaris (version 7.2.1) software (Bitplane). For ease of EOM visualization, the Z-stack volumes were first manually segmented to define the EOM or the whole POM area using the Isosurface Imaris function. The signal outside the isosurface was set to zero, the corresponding channel duplicated and subsequently a new isosurface was created using automatic thresholding on the new channel.

### In Situ Hybridization

Whole-mount in situ hybridization with digoxigenin-labeled antisense mRNA probes was performed as described previously (Comai et al., 2014). The Aldh1a1, Aldh1a2, Aldh1a3 probes were previously described (Matt et al., 2005; Molotkov et al., 2006).

### micro-CT (micro computed tomography) analysis

The tissue contrasting protocol has been adapted from the original protocol developed by (Metscher, 2009) and applied to mouse embryos as described in (Kaucka et al., 2018) and (Heude et al., 2018). For tissue contrasting, E13.5 embryos were stained in 0.5% PTA (phospho-tungstic acid) in 90% Methanol for 4 days, E15.5 embryos were stained in 0.7% PTA in 90% Methanol for 1 week.

The μCT analysis of the embryos was conducted using the GE phoenix v|tome|x L 240 (GE Sensing and Inspection Technologies GmbH, Germany), equipped with a 180 kV/15W maximum power nanofocus X-ray tube and flat panel detector DXR250 with 2048 × 2048 pixel, 200 × 200 μm pixel size. The exposure time of the detector was 900 ms in every position over 360°. Three projections were acquired and averaged in every position for reduction of the noise in μCT data. The acceleration voltage was set to 60 kV and tube current to 200 μA. The radiation was filtered by 0.2 mm of aluminium plate. The voxel size of obtained volumes (depending on a size of an embryo) appeared in the range of 2 μm - 6 μm. The tomographic reconstructions were performed using GE phoenix datos|x 2.0 3D computed tomography software (GE Sensing and Inspection Technologies GmbH, Germany). The EOM, eye and cartilages in the embryo head were segmented by an operator with semi-automatic tools within Avizo - 3D image data processing software (FEI, USA). The 3D segmented region was transformed to a polygonal mesh as a STL file, imported to VG Studio MAX 2.2 software (Volume Graphics GmbH, Germany) for surface smoothing and 3D visualisation.

### Cell isolation from the periocular region

The periocular region of *Rare^CreERT2^:R26^mTmG^* embryos (including the eye itself) was micro-dissected and minced with small scissors inside a 2ml Epperndorf tube. 1ml of TrypLE™ Express (Invitrogen, 12604013) was added to each tube incubated for 15min at 37°C with agitation. Samples were resuspended by gently pipetting up and down 10-15 times using a P1000 pipette. Upon addition of 1ml of culture media containing (10µg/ml) DNaseI (Roche, 11284932001), samples were spun 15’ 500g at RT, pellet resuspended in 400ul of culture media containing of 20% foetal bovine serum (FBS; Gibco), 1% Penicillin-Streptomycin (15140 Gibco), 2% Ultroser G (15950-017, Pall Biosepra) in 50:50 DMEM:F12 (31966 and 31765, Gibco) and plated on individual wells of 8 well glass bottom dishes (Ibidi, 80826) coated with 1 mg/ml of Matrigel (354234, BD Biosciences). Cells were allowed to attach for 8h at 37 °C 5% CO_2_, washed with PBS and fixed 15 min RT with 4% PFA/PBS. After fixation, cells were washed in PBS and permeabilized with 0.5% Triton in PBS X-100 5 min at RT. After three washes in PBS (5min each), cells were blocked with 20% goat serum in PBS 1 h at RT. Primary antibodies were added to cells in 2% goat serum in PBS for 2 h at RT or ON at 4C. Cells were washed three times with PBS, incubated with secondary antibodies 1 h at RT, washed in PBS and kept in PBS for imaging.

### Statistics

The number of embryos of each genotype used for analysis is indicated in the figure legends and Table 1. The graphs were plotted and statistical analyses were performed using Prism8 (GraphPad Software, Inc). All data points are presented as mean ± SEM (error bars). Statistical tests used for analysis are indicated on the respective figure legends. P values less than 0.05 were considered significant (*p < 0.05; **p < 0.01; ***p < 0.001).

## Acknowledgements

We acknowledge funding support from the Institut Pasteur, Association Française contre le Myopathies, Agence Nationale de la Recherche (Laboratoire d’Excellence Revive, Investissement d’Avenir; ANR-10-LABX-73) and the Centre National de la Recherche Scientifique. We gratefully acknowledge the UtechS Photonic BioImaging (Imagopole), C2RT, Institut Pasteur, supported by the French National Research Agency (France BioImaging; ANR-10–INSB–04; Investments for the Future). MT, TZ and JK acknowledge the project CEITEC 2020 (LQ1601) with financial support from the Ministry of Education, Youth and Sports of the Czech Republic under the National Sustainability Programme II and Ceitec Nano+ project CZ.02.01/0.0./.0.0./16_013/0001728 under the program OP RDE. MT was financially supported by by the Brno City Municipality as a Brno Ph.D. Talent Scholarship Holder. The funders had no role in study design, data collection and analysis, decision to publish, or preparation of the manuscript.

